# Aerobic methoxydotrophy: growth on methoxylated aromatic compounds by *Methylobacterium*

**DOI:** 10.1101/712836

**Authors:** Jessica A. Lee, Sergey Stolyar, Christopher J. Marx

## Abstract

Pink-pigmented facultative methylotrophs have long been studied for their ability to grow on reduced single-carbon (C1) compounds. The C1 groups that support methylotrophic growth may come from a variety of sources. Here, we describe a group of *Methylobacterium* strains that can engage in methoxydotrophy: they can metabolize the methoxy groups from several aromatic compounds that are commonly the product of lignin depolymerization. In addition, these organisms can utilize the full aromatic ring as a growth substrate, a phenotype that has rarely been described in *Methylobacterium*. We demonstrated growth on *p*-hydroxybenzoate, protocatechuate, vanillate, and ferulate in laboratory culture conditions. We also used comparative genomics to explore the evolutionary history of this trait, finding that the capacity for aromatic catabolism is likely ancestral to two clades of *Methylobacterium*, but has also been acquired horizontally by closely related organisms. In addition, we surveyed the published metagenome data to find that the most abundant group of aromatic-degrading *Methylobacterium* in the environment is likely the group related to *M. nodulans*, and they are especially common in soil and root environments. The demethoxylation of lignin-derived aromatic monomers in aerobic environments releases formaldehyde, a metabolite that is a potent cellular toxin but that is also a growth substrate for methylotrophs. We found that, whereas some known lignin-degrading organisms excrete formaldehyde as a byproduct during growth on vanillate, *Methylobacterium* do not. This observation is especially relevant to our understanding of the ecology and the bioengineering of lignin degradation.

## 1 Introduction

Microbial processes for degrading lignin and lignin-derived aromatics are of intense interest in a diversity of fields, ranging from fundamental ecosystems science—which seeks to understand the processes by which the carbon fixed in plant biomass is mineralized (Lehmann and Kleber, 2015)—to bioenergy engineering—where the recalcitrance of lignin is a major hurdle in the processing of plant biomass (Ragauskas et al., 2014)—and the fossil fuel industry—which seeks to understand microbial transformations of coal (Welte, 2016). Lignin, which comprises approximately 20% of the carbon fixed by photosynthesis on land (Ruiz-Dueñas and Martínez, 2009), is an exceptionally complex, irregular, polycyclic aromatic polymer, in which many of the constituent aromatic rings are heavily substituted with methoxy (-OCH_3_) groups (Vanholme et al., 2010). While depolymerization of lignin in the environment is typically attributed to the activity of fungi, recent studies probing the biodiversity of the microbial lignin-degrading community in soils have inspired increased interest in the role of bacteria in both depolymerization and in the degradation of its monomeric aromatic constituents (Wilhelm et al., 2019; Díaz-García et al., 2021).

In the aerobic bacterial degradation of lignin-derived aromatic monomers such as vanillate, the degradation of the aromatic ring proceeds by the protocatechuate branch of the beta-ketoadipate pathway. The gene cluster encoding this pathway is widely distributed among soil microorganisms, and has a complex evolutionary history resulting in diverse patterns of gene organization and regulation (Harwood and Parales, 1996; Parke, 1997; Buchan et al., 2004). In the case of methoxylated aromatics, ring cleavage is preceded by the removal of the methoxy group; in most aerobic organisms, this occurs via vanillate monooxygenase, a Rieske [2Fe-2S] enzyme that demethylates vanillate to generate protocatechuate and formaldehyde (Brunel and Davison, 1988; Mitsui et al., 2003; Merkens et al., 2005; Sudtachat et al., 2009; Chen et al., 2012) (Fig. 1). *Pseudomonas putida* growing on vanillate as a sole carbon source excretes formaldehyde into the medium in measurable amounts (Lee et al., 2021). Formaldehyde is a potent electrophile, and toxic to microorganisms due to its reactivity with DNA and proteins (Chen et al., 2016). Elimination of this toxin is therefore essential to lignin degradation. Multiple studies have demonstrated experimentally, through either engineered disruption or constitutive expression of formaldehyde oxidation capacity in *Bradyrhizobium diazoefficiens* (Sudtachat et al., 2009), *Pseudomonas putida* (Hibi et al., 2005), *Burkholderia cepacia* (Mitsui et al., 2003), and *Corynebacterium glutamicum* (Lessmeier et al., 2013), that demethoxylation of vanillate is dependent on the activity of a functional formaldehyde detoxification system, and formaldehyde removal may be the rate-limiting step to the degradation of lignin-derived methoxylated aromatics.

**Figure 1.**
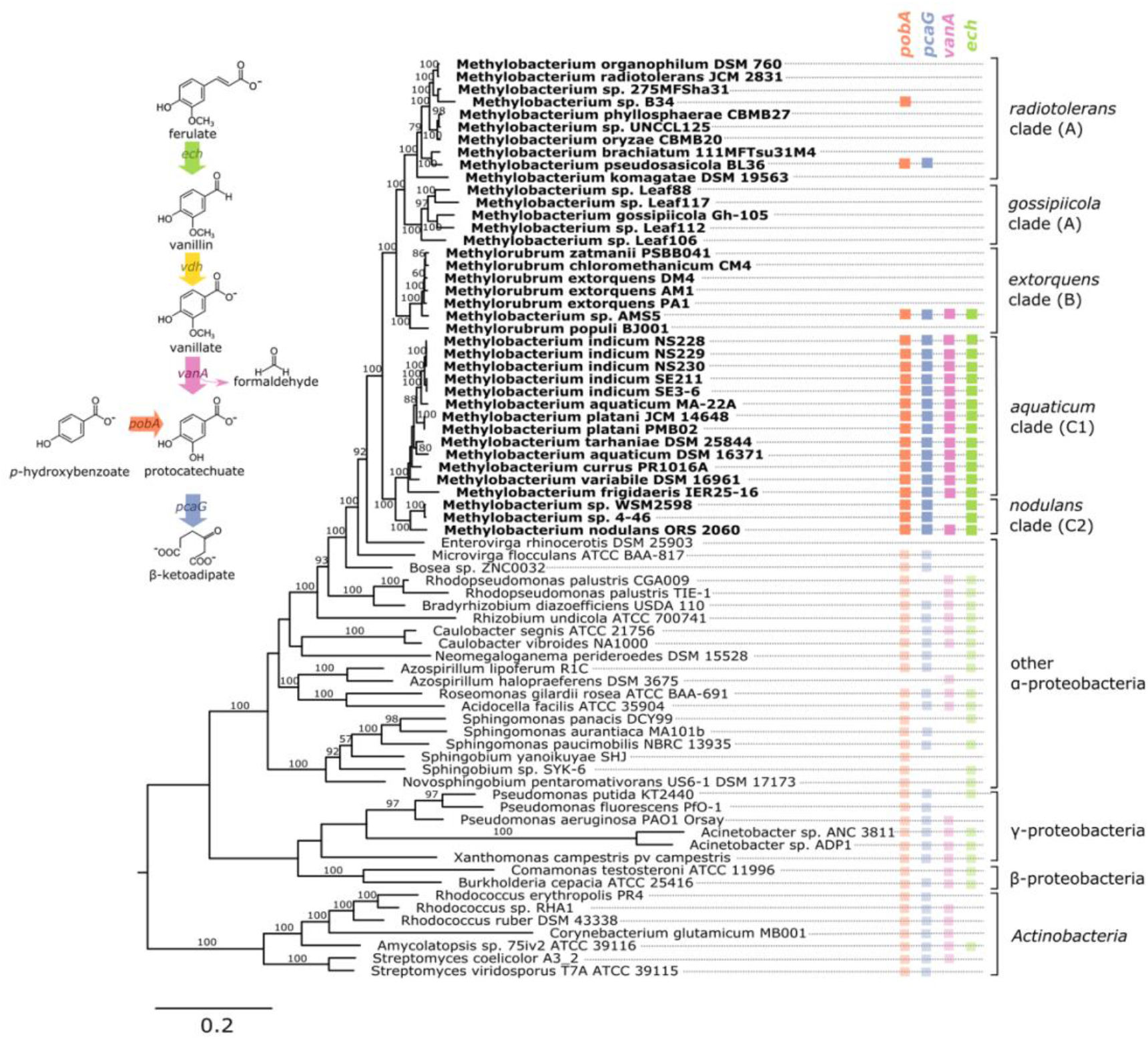
Genes associated with aromatic catabolism are present in two clusters of closely-related *Methylobacterium* species, with some exceptions. Genomes of *Methylobacterium* species and known aromatic-degrading bacteria were searched for four genes involved in different steps of the degradation of lignin-derived methoxylated aromatic compounds (upper left). Colored squares indicate the presence of each gene. *Methylobacterium* species are shown in bold text and colors, reference organisms (not *Methylobacterium*) in normal text and lighter colors. Note: among the reference organisms, all species that do not have *pcaG* do have the genes for protocatechuate 4,5-dioxygenase, indicating an alternative ring cleavage mechanism and different pathway for the catabolism of PCA. The phylogeny was composed using a concatenated alignment of four housekeeping genes (*atpD, recA, rpob*, and 16S rRNA), with *Prochlorococcus marinus* as an outgroup (not shown). Clade A/B/C designations are in accordance with (Green and Ardley, 2018; Alessa et al., 2021; Leducq et al., 2021). Branch labels indicate bootstrap percent; values <50 are not shown. Sequence data are given in Supplementary Data Files 1-4. Due to space considerations, not all genomes searched are shown in this figure; the full set, with accession numbers, is given in Supplementary Tables 1 (*Methylobacterium*) and 2 (reference species), and a phylogeny of all *Methylobacterium* species are shown in Supplementary Figure 2.

Formaldehyde detoxification pathways in lignin-degrading organisms are diverse, and in some organisms include more than one mechanism. In some cases formaldehyde is oxidized to CO2 via formate, such as in the thiol-dependent systems in *C. glutamicum* and *B. japonicum* or the Zn-dependent aldehyde dehydrogenase in *P. putida* (Roca et al., 2009); in others it may be incorporated into biomass via the ribulose monophosphate (RuMP) pathway, as in *B. cepacia* (Kato et al., 2006). Furthermore, methanogens of the genus *Methermicoccus* are capable of using the methoxy groups of coal-derived aromatic compounds for the production of methane through a novel metabolic pathway (“methoxydotrophic methanogenesis”), a discovery that illuminated a previously-unknown link between C1 metabolism and the anaerobic degradation of recalcitrant aromatic compounds (Mayumi et al., 2016; Lloyd et al., 2021).

Methylotrophs, obligately aerobic organisms that use reduced C1 compounds for growth, therefore make appealing candidates for efficient degradation of lignin-derived methyoxylated aromatics. Members of the pink-pigmented methylotrophic alpha-proteobacteria, and particularly the model organism *Methylorubrum* (formerly *Methylobacterium) extorquens*, have long been studied for their metabolism of simple C1 compounds such as methanol, formate, methylamine, and halogenated methanes (Chistoserdova, 2011). Formaldehyde is a central intermediate in the metabolism of many of these substrates, raising the possibility that, if these methylotrophs were capable of demethoxylating aromatic compounds, they could also use the resulting formaldehyde as a growth substrate (Lee et al., 2021). It has previously been documented that two members of the group are capable of growth on aromatic compounds (Ito and Iizuka, 1971; Jourand, 2004; Ventorino et al., 2014); however, the prevalence of this trait among *Methylorubrum, Methylobacterium*, and their close relatives, and the organization and evolutionary history of the genes involved, has not been described. And while formaldehyde metabolism has been extensively studied in the pink methylotrophs (Vorholt et al., 1998; Marx et al., 2003), aromatic compounds have traditionally not been discussed as a potential substrate for methylotrophic growth (Chistoserdova and Lidstrom, 2013; Kelly et al., 2014).

Here we report novel findings on the ecology and evolution of methoxydotrophic growth and catabolism of aromatic compounds by members of the *Methylobacterium*. We explored the genomic capacity of these species by searching published genomes, verified growth and formaldehyde production/consumption on aromatic compounds in the laboratory, and surveyed published metagenome data to assess the distribution and prevalence of aromatic-degrading *Methylobacterium* in the environment. For a thorough description of aerobic microbial degradation of lignin-derived aromatic compounds, we direct the reader to the excellent reviews published previously (Harwood and Parales, 1996; Masai et al., 2007); a simplified diagram of the pathway and the four genes of interest to this study (*vanA, pcaG, pobA, ech*) is given in Figure 1. Because we were especially interested in the interaction between aromatic catabolism and methylotrophy, our metagenomic analysis focused especially on *vanA* (KO:K03862), the gene encoding the alpha subunit of vanillate monooxygenase.

## 2 Materials and Methods

### 2.1 Taxonomic nomenclature

A recently proposed re-organization of the taxonomy of the pink-pigmented facultative methylotrophs suggested renaming the *extorquens* clade with the genus name “*Methylorubrum*” (Green and Ardley, 2018). However, others have pointed out that the remaining *Methylobacterium* clades are not monophyletic, leading to the suggestion that the entire group be returned to the name *Methylobacterium*, until more data is gathered to support splitting the genus (Hördt et al., 2020; Alessa et al., 2021). Therefore, in this manuscript, we will refer to the formally described *Methylorubrum* species using that name, but to describe the larger monophyletic group that is the focus of this study we will use the term *Methylobacterium*.

### 2.2 Phylogenetic analysis

The genetic potential of *Methylobacterium* species (Kelly et al., 2014) (Table S1) for the degradation of aromatic compounds was assessed through analysis of genomes available on the IMG/M database (Chen et al., 2019). We surveyed all the genomes that were classified in the genera *Methylobacterium* or *Methylorubrum* as of April 2020. Of the 134 genomes returned, 3 were omitted from analysis: *Methylobacterium* sp. CCH7-A2 is classified by the Genome Taxonomy Database (Parks et al., 2018) as a *Bosea* sp. and by IMG as forming a clique with *Porphyrobacter donghaensis*; *Methylobacterium* sp. ZNC0032 is also annotated as a *Bosea* sp. and does not cluster with the *Methylobacterium* (included in Fig. 1 and Supplementary Table 1 but not in further analyses); and *Methylobacterium platani* JCM 14648 has 2 genome entries in IMG, of which we used the earlier, more complete assembly. We should note that we found that a genome currently named *Streptomyces purpurogeneiscleroticus* NRRL B-2952 is identified as *Methylobacterium* by GTDB and in IMG forms a clique with several *Methylobacterium* species, including *M. fujisawaense*, *M. oryzae*, and *M. phyllosphaerae*; however, it does not carry the *vanA* gene and was not included in this analysis. Finally, it has been noted that the published genome of *M. organophilum* used in this study may contain contaminants and a resequencing places it in closer relation to the *extorquens* (B) clade than shown here (Alessa et al., 2021), but it is not one of the *vanA*-carrying organisms of interest. In total we analyzed 131 genomes, described in Supplementary Table 1.

Genes of interest were identified by Kyoto Encyclopedia of Genes and Genomes (KEGG) Orthology ID: *vanA* (vanillate monooxygenase alpha subunit, KO:K03862); *pcaG* (protocatechuate 3,4-dioxygenase alpha subunit, KO:K00448); *pobA* (*p*-hydroxybenzoate 3-monooxygenase, KO:K00481); and *ech* (*trans*-feruloyl-CoA hydratase/vanillin synthase, KO:K18383). To identify ech genes, we also searched for IMG Term Object ID 8865, vanillin synthase, and we used IMG’s Conserved Neighborhood search tool to identify an ech homolog in that was syntenic across several *Methylobacterium* genomes but only sometimes annotated as Vanillin Synthase). Locus tags of all genes are given in Supplementary Table 1. Vanillin synthase, *vdh* (KO:K21802), was not studied because we were not easily able to identify homologs in all organisms. Known lignin-degrading organisms were added to the phylogenies as reference organisms, and *Prochlorococcus marinus* NATL2A and archaeal species were used as outgroups (Supplementary Table 2).

For the phylogeny in Figure 1, which shows all putative aromatic-degrading *Methylobacterium* strains in the context of selected other *Methylobacterium* and known lignin-degrading organisms, four gene alignments were concatenated: 16S rRNA (the RNA component of the small ribosomal subunit), *rpoB* (DNA-directed RNA polymerase subunit beta, KO:K03043), *atpD* (ATP Synthase F1 complex beta subunit, KO:K02112), and *recA* (recombination protein RecA, KO:K03553), all of which have been shown previously to be informative for phylogenetic analysis in *Methylobacterium* and closely related organisms (Gaunt et al., 2001; Menna et al., 2009; Zhang et al., 2012). Locus tags are given in Supplementary Table 1. The clade structure generally agrees with previous *Methylobacterium* phylogenies based on 16S rRNA, *rpoB*, or core genes, though some analyses find clade A not to be monophyletic (Green and Ardley, 2018; Alessa et al., 2021; Leducq et al., 2021).

Two additional phylogenies were constructed as supplementary information: one containing all the *Methylobacterium* strains surveyed for this study, using the *rpoB* gene (Leducq et al., 2021) with *Bradyrhizobium diazoefficiens* USDA 110 as an outgroup; and one showing the relationships of the Rieske [2Fe-2S] oxygenase homologs. The *rpoB* tree includes five of the *Methylobacterium* strains carrying aromatic catabolism genes (sp. MIMD6, sp. 174MFSha1-1, sp. AM11, sp. YR596, and sp. CCH7-A2) for which the full-length 16S rRNA gene sequence was not available and therefore are not included in Fig. 1. For 3 of the *Methylobacterium* strains in our study (*Methylobacterium* sp. CG08_land_8_20_14_0_20_71_15, *Methylobacterium phyllosphaerae* JCM 16408, and *Methylorubrum thiocyanatum* JCM 10893), no *rpoB* sequences were available and aromatic catabolism genes were also not detected; these strains do not appear in any of the phylogenies but are listed in Supplementary Table 1. The homolog tree was constructed using all the genes annotated as COG 4638 in IMG in the surveyed *Methylobacterium* strains, as well as *vanA* sequences from other species for reference.

All gene sequences were aligned using MUSCLE v. 3.7 (Edgar, 2004) (RRID:SCR_011812) on the CIPRES Science Gateway (Miller et al., 2010), using the following parameters at all iterations: ClustalW as the sequence weighting scheme, UPGMB clustering, SUEFF=0.1, pseudo rooting, and sum-of-pairs scoring for refinement. Phylogenetic trees were generated using RAxML v. 8.2.12 (Stamatakis, 2014) (RRID:SCR_006086) on CIPRES using the RAxML-HPC2 on XSEDE tool, with the following parameters: 25 rate categories, rapid bootstrapping with GTRCAT model for 100 iterations, random seed=12345 for bootstrapping and parsimony, and outgroups as given in Table S2 (outgroups are not shown in figures). All alignments are provided in Supplementary Data Sheets 1-10.

### 2.3 Genomic context of aromatic catabolism genes

We used GeneHood v.0.15.0-0 (Ortega, 2020) to assess synteny in the gene neighborhoods surrounding the *vanA* and *pcaH* genes in *Methylobacterium* species. For each gene of interest, the database locus tag from the MIST 3.0 database (Gumerov et al., 2020) was provided and up to 15 genes upstream and downstream of the gene were plotted (in genomes that are not fully assembled, some scaffolds were did not have 15 genes on either side). Phylogenetic trees, gene annotations, and color-coding were added manually using Inkscape v. 0.91 (Inkscape Team, 2003) (RRID:SCR_014479).

GC content in the *Methylobacterium* sp. AMS5 genome was calculated in R v.4.0.2 (R Core Team, 2018) (RRID:SCR_001905) using the alphabetFrequency function from Biostrings v.2.46.0 (Pagès et al., 2019) (RRID:SCR_016949); for plotting, a sliding window of 5 kb was used for the full genome and 500 bp for the catabolic island region. Genome signature difference was calculated using a custom script in R, using the formula for the δ* difference (Karlin, 2001) with tetranucleotides rather than dinucleotides.

### 2.4 Growth assays on aromatic substrates

All organisms were grown on MPIPES medium (Delaney et al., 2013) with the addition of 1x Wolfe’s Vitamins (Atlas, 2010), 25 μM LaCl_3_ (as has been found to facilitate growth in some *Methylobacterium* spp. (Skovran and Martinez-Gomez, 2015)), and carbon substrates as specified. All conditions contained 16 mM of carbon (16 mM methanol, 4 mM succinate, 2.67 mM glucose, 2.29 mM benzoate, 2.29 mM *p*-hydroxybenzoate (PHBA), 2.29 mM protocatechuic acid (PCA), 2 mM vanillate, 1.6 mM ferulate, or 0.4 g/L Kraft lignin). While some soil microenvironments may experience higher substrate concentrations and some lower, 16 mM of carbon is in the range that has been commonly used for *M. extorquens* to ensure that final population size scales linearly with substrate concentration for both single- and multi-carbon substrates (Lee et al., 2009). *Methylobacterium* sp. 4-46 was also tested on methanol without LaCl_3_ to assess the effect of lanthanum on methanol growth. For each assay, one colony was inoculated from a culture plate into 5 mL MPIPES with 3.5 mM disodium succinate and grown 24 hours to obtain a stationary-phase culture, then diluted 1:64 (v/v) in the same medium and grown again until stationary. Using that inoculum, the growth experiment was conducted as follows: each replicate was diluted 1:64 into MPIPES and aliquoted into several wells of a Costar 48-well tissue culture-treated plate (product #3548, Corning Inc., Corning, NY) for non-volatile substrates, or into Balch-type glass culture tubes for methanol, along with 16 mM carbon substrate. Total volume was 640 μL per well in culture plates or 5 mL per tube in Balch tubes. Each strain was tested in biological triplicate (3 separate colonies). Balch tubes were sealed with butyl rubber stoppers to prevent escape of methanol. Plates were incubated at 30 °C in a LPX44 Plate Hotel (LiCONiC, Woburn, MA) with shaking at 250 RPM. Optical density was assessed using a Wallac 1420 Victor2 Microplate Reader (Perkin Elmer, Waltham, MA), reading OD_600_ for 0.4 seconds at intervals of between 2 and 5 hours. Culture tubes were incubated in a TC-7 tissue culture roller drum (New Brunswick Scientific, Edison, NJ) at a speed setting of 7, and OD_600_ was measured in the culture tubes using a Spectronic 200 spectrophotometer (Thermo Fisher Scientific, Waltham, MA). Outputs from the Wallac 1420 software were collated using Curve Fitter (Delaney et al., 2013); data cleaning and plotting were then conducted in R.

### 2.5 Formaldehyde production during growth on vanillate

Growth of *Methylobacterium* strains on vanillate was initiated with stationary-phase MPIPES-succinate cultures as described above; these were diluted 1:64 (v/v) into 5 mL MPIPES with 2 mM vanillate and grown 2 days until stationary phase to allow cultures to acclimate to the substrate. This inoculum was diluted 1:32 into culture flasks containing 20 mL of MPIPES with 2 mM vanillate and incubated, shaking, at 30 °C. Cultures were sampled every 4 hours: OD_600_ was measured in a SmartSpec Plus Spectrophotometer (Bio-Rad Laboratories, Hercules, CA), and the supernatant of 500 μL of centrifuged culture was used for the measurement of formaldehyde using the colorimetric assay described by Nash (Nash, 1953). All strains were tested in biological triplicate. Although the active component in the Nash assay used has been found to react to several multi-carbon aldehydes, we have no reason to believe these compounds could be produced by vanillate metabolism, given what is known about the pathway; moreover, reactions with all compounds are slower than the reaction with formaldehyde (Compton and Purdy, 1980).

Formaldehyde production of non-methylotrophs during vanillate growth was assayed similarly. Stationary-phase vanillate-grown cultures of *Pseudomonas putida* KT2440 and *Rhodococcus jostii* RHA1 (USDA Agricultural Research Service Culture Collection (NRRL)) were diluted 1:64 (v/v) into 5 mL MPIPES medium in culture tubes, with sampling every 2 hours. Vanillate was provided at 4 mM; although this was greater than in the *Methylobacterium* experiment, we have found that vanillate concentration has little effect on the generation of formaldehyde in the medium (Lee et al., 2021).

### 2.6 Analysis of *Methylobacterium vanA* genes in published metagenomes

To assess the abundance of *Methylobacterium*-encoded *vanA* genes in the environment, we searched all published metagenomes available through the IMG/M portal on December 20, 2017. The “Function Search” tool was used to identify all metagenomes containing genes annotated with KO:K03862 (*vanA*). To increase our chances of observing meaningful ecological patterns, we restricted our analysis to metagenomes containing >100 *vanA* genes. Genes on scaffolds with a phylogenetic lineage assignment of *Methylobacterium* were designated “*Methylobacterium vanA*.” and the Ecosystem Type field in the sample metadata was summarized into a “Sample Type” designation for analysis (e.g., Phyllosphere and Phylloplane were classified as “Leaf,” Rhizosphere and Rhizoplane as “Root,” etc).

To provide a tree-based assessment of *vanA* diversity in addition to the USEARCH-based IMG/M phylogenetic lineage assignment (Huntemann et al., 2016), we downloaded the *Methylobacterium vanA* gene sequences and placed them in the reference *vanA* phylogeny (described above). pplacer v.1.1 (Matsen et al., 2010) (RRID:SCR_004737) was used with HMMer v.3.1b.2 (Eddy, 2007) (RRID:SCR_005305), RAxML v.8.2 (Stamatakis, 2014), and taxtastic v.0.8.8 for Python 3 (RRID:SCR_008394). The resulting tree was paired with metadata from sequence scaffolds in Phyloseq v.1.22.3 for R (McMurdie and Holmes, 2013) (RRID:SCR_013080) for the final phylogeny. Using pplacer, we found that only 182 out of 348 genes actually fell within the *Methylobacterium* clades, whereas 31 clustered more closely with *Azospirillum halopraeferens*, and the remaining 135 were distributed among more distantly related organisms (Supplementary Figure 1). This may be due in part to the overall complexity of the Rieske-type monooxygenase family (Özgen and Schmidt, 2019). We therefore took the conservative approach of focusing only on the genes identified by both IMG and pplacer as *Methylobacterium* for the remainder of our analyses.

A separate analysis was carried out to assess the relative abundances of all *Methylobacterium* reads within each metagenome in the JGI IMG database. The “Phylogenetic Distribution: Genomes v. Metagenomes” tool was used; this tool conducts a BLAST search to find reads in a set of metagenomes that have similarity to a query genome (JGI IMG, 2017). 18 query genomes were used: *M. radiotolerans* JCM 2831, *M. pseudosasiacola* BL36, *M. extorquens* PA1, *Methylobacterium sp*. AMS5, *Methylobacterium sp*. Leaf88, *M. gosipiicola* Gh-105, *M. indicum* SE3.6, *M. indicum* SE2.11, *M. aquaticum* MA-22A, *M. platani* JCM 14648, *M. variabile* 16961, *M. tarhaniae* DSM 25844, *M. aquaticum* DSM 16371, *Methylobacterium sp*. WSM2598, *Methylobacterium sp*. 4-46, *M. nodulans* ORS 2060, *Microvirga flocculans* ATCC BAA-817, and *Enterovirga rhinocerotis* DSM 25903 (genome accessions in Supplementary Table 1). A total of 33,967 metagenomes were searched; this search set comprised all metagenomes published in IMG as of January 10, 2021 that were returned by the following 6 search terms:

- GOLD Ecosystem = Host-Associated & GOLD Ecosystem Category = Plant
- GOLD Ecosystem = Host-Associated & GOLD Ecosystem Category = Human
- GOLD Ecosystem = Host-Associated & GOLD Ecosystem Category = Mammals
- GOLD Ecosystem = Environmental & GOLD Ecosystem Category = Terrestrial
- GOLD Ecosystem = Environmental & GOLD Ecosystem Category = Aquatic
- GOLD Ecosystem = Engineered

Metagenomes with a published gene count of 0 were excluded from analysis, but otherwise no attempt was made to sample evenly across ecosystems, or to screen metagenomes by size, assembly status, etc., as the goal was to be as complete as possible in the search of available data. Due to computational constraints of IMG, metagenomes were searched in groups of 5,000 or fewer and 1 genome at a time, and results were combined. The output of each search consisted of a data table including identity (and metadata) of each metagenome sample where the genome was found, and the number of reads that hit to the genome at the 90% similarity level.

## 3 Results

### 3.1 Genetic capacity for metabolism of methoxylated aromatic compounds is present primarily in two clades of *Methylobacterium*

To investigate the possibility of aromatic catabolism among *Methylobacterium* strains, we first searched 131 published genomes for genes encoding four key enzymes known to be involved in the degradation pathway of methoxylated aromatic compounds: *vanA* (vanillate monooxygenase alpha subunit, KO:K03862);*pcaG* (protocatechuate 3,4-dioxygenase alpha subunit, KO:K00448); *pobA* (*p*-hydroxybenzoate 3-monooxygenase, KO:K00481); and *ech* (*trans*-feruloyl-CoA hydratase/vanillin synthase, KO:K18383; we searched also for IMG Term Object ID 8865, vanillin synthase, and used genome synteny; see Methods). We found that all four genes are indeed present within the genus *Methylobacterium*. They are primarily limited to two closely related clades, the *aquaticum* clade (C1) and the *nodulans* clade (C2); furthermore, they are present in nearly all the genomes we surveyed in these clades (Fig. 1, Supplementary Table 1, Supplementary Figure 2). However, there are exceptions. Of the genomes searched, *Methylobacterium sp*. AMS5 in the *extorquens* clade (clade B) and *M. pseudosasiacola* in the *radiotolerans* clade (clade A) also carry some aromatic catabolism genes (Fig. 1).

Two members of clade C2 (sp. 4-46 and sp. WSM2598) appear to lack *vanA* but carry *ech* (Fig. 1). It is possible that the *ech* homologs annotated as *trans*-feruloyl-CoA hydratase/vanillin synthase in fact carry out a different reaction in these organisms (Lohans et al., 2017); however, they are located adjacent other genes annotated as vanillin dehydrogenase (see below), which catalyzes the conversion of vanillin to vanillate. This suggests that *Methylobacterium* sp. 4-46 and sp. WSM2598 may have previously had the capacity to utilize the product of these reactions but has since lost *vanA*. Among the closest sister taxa to *Methylobacterium, Enterovirga rhinocerotis* has none of the four genes of interest, and *Microvirga flocculans* has only *pobA* and *pcaH*.

While the absence of a gene in a draft-status genome is not always an indication that the gene is not there, several of the genomes without aromatic catabolism genes, including *M*. sp. 4-46 and *M. extorquens* AM1 and PA1, are complete (Vuilleumier et al., 2009; Marx et al., 2012) (Supplementary Table 1). And the substantial number of genomes we observed without any evidence of aromatic catabolism genes suggested a broad phylogenetic pattern: the capacity to metabolize several lignin-derived aromatic compounds was likely acquired by clade C early in its divergence within the *Methylobacterium*, but members of other clades have acquired the genes more recently by horizontal gene transfer. We sought further support for this evolutionary hypothesis by analyzing the phylogenies of the genes themselves, examining the genomic contexts of *vanA* and *pcaG*, and assaying the ability of several strains to grow on methoxylated aromatic compounds in the lab.

### 3.2 Phylogeny of aromatic catabolism genes supports ancestral origins in *M. aquaticum* and *M. nodulans* clades and horizontal acquisition by two other species

The phylogenies of *pobA, pcaG*, and *vanA* were largely congruent with the phylogeny of conserved housekeeping genes within the *Methylobacterium* but not between *Methylobacterium* and their closest relatives, supporting the hypothesis that the four genes are ancestral to clade C but were not necessarily acquired together (Fig. 2). Phylogeny also supported our hypothesis of the function of the *vanA* gene in *Methylobacterium* species: although *vanA* homologs, members of the Rieske [2Fe-2S] oxygenases, are known to carry out a variety of reactions (Kweon et al., 2008; Özgen and Schmidt, 2019), and *Methylobacterium* species have several such genes, only the genes annotated as *vanA* cluster with genes that have been characterized as vanillate monooxygenases in known lignin-degrading bacteria (Supplementary Figure 3). The *ech* phylogeny was the most difficult to interpret, as the *Methylobacterium* separated into three clades (Fig. 2). It is possible that not all gene homologs in the phylogeny code for enzymes specific to ferulate; further biochemical and genetic work will be necessary to be certain. Notably, *Methylobacterium* sp. AMS5 and *M. pseudosasiacola* were more closely related to each other than to any other species in both their *pobA* and *pcaG* sequences, despite their phylogenetic distance from each other at the genome level. And while the *pobA* sequences of those two strains fell within those of other *Methylobacterium*, their *pcaG* sequences clustered with the *Sphingomonas*, further support not only for the hypothesis that these two disparate species both acquired the beta-ketoadipate pathway via horizontal gene transfer, but that they may have acquired it from the same donor or that one served as the donor to the other.

**Figure 2.**
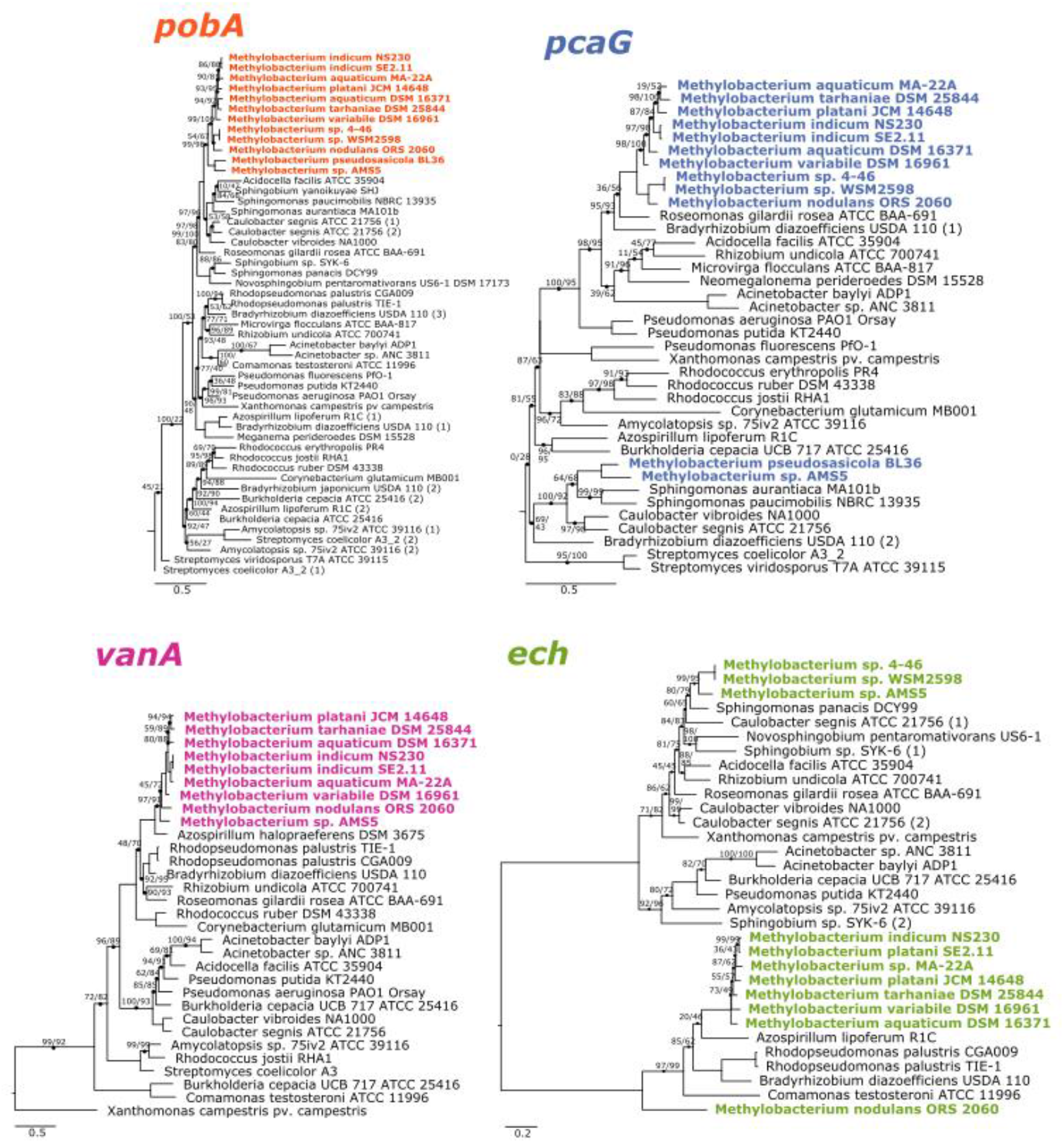
Phylogenies of aromatic catabolism genes suggest that all four are ancestral to the *M. aquaticum/nodulans* clades and have been horizontally acquired by two other *Methylobacterium* species. For all organisms in Fig. 1, the sequences of *pobA* (*p*-hydroxybenzoate 3-monooxygenase), *pcaH* (protocatechuate 3,4-dioxygenase beta subunit), *vanA* (vanillate monooxygenase subunit A), and *ech* (*trans*-feruloyl-CoA hydratase/vanillin synthase) were aligned and phylogenies constructed using maximum likelihood, with homologs from archaeal organisms as outgroups (not shown). Accession numbers for all genes are given in Tables S1 and S2. Branch labels indicate bootstrap support. Comparison of these phylogenies and that in Fig. 1 suggest that all four genes are restricted to and prevalent in the *M. aquaticum* and *M. nodulans* clades, with a few losses (*vanA* by *Methylobacterium sp*. 4-46 and WSM2598) and some gains by horizontal gene transfer (in *M. pseudosasiacola* and *Methylobacterium sp*. AMS-5).

### 3.3 Gene synteny in aromatic catabolism gene neighborhoods is consistent with phylogeny

We further examined the relationships of *vanA* and *pcaG* across species by comparing patterns of synteny within the genome neighborhood for each (Fig. 3). In all strains with *pcaG*, the gene was located within a cluster of beta-ketoadipate pathway genes, and five of these genes were present in all strains in the same order. Strikingly, we found no genes annotated as *pcaI* and *pcaJ* (beta-ketoadipate:succinyl-CoA transferase) or *pcaF* (encoding beta-ketoadipyl-CoA thiolase) within these operons, which are necessary for the final steps of the beta-ketoadipate pathway (Harwood and Parales, 1996; Buchan et al., 2004), leaving it initially unclear whether and how these organisms might be able to metabolize vanillic acid and related compounds. However, almost all strains contained an additional gene between *pcaG* and *pcaB*, annotated as an uncharacterized conserved protein DUF849 (COG3246, beta-keto acid cleavage enzyme). In all *Methylobacterium* strains where *vanA* was found, it was accompanied by the other two genes required to confer the ability for the demethoxylation of vanillate: *vanB* (encoding the other subunit of vanillate monooxygenase) and *vanR* (an AraC family transcriptional regulator). Some strains also had genes encoding a Major Facilitator Superfamily transporter specific to protocatechuate (*pcaK*) or vanillate (*vanK*) (Fig. 3).

**Figure 3.**
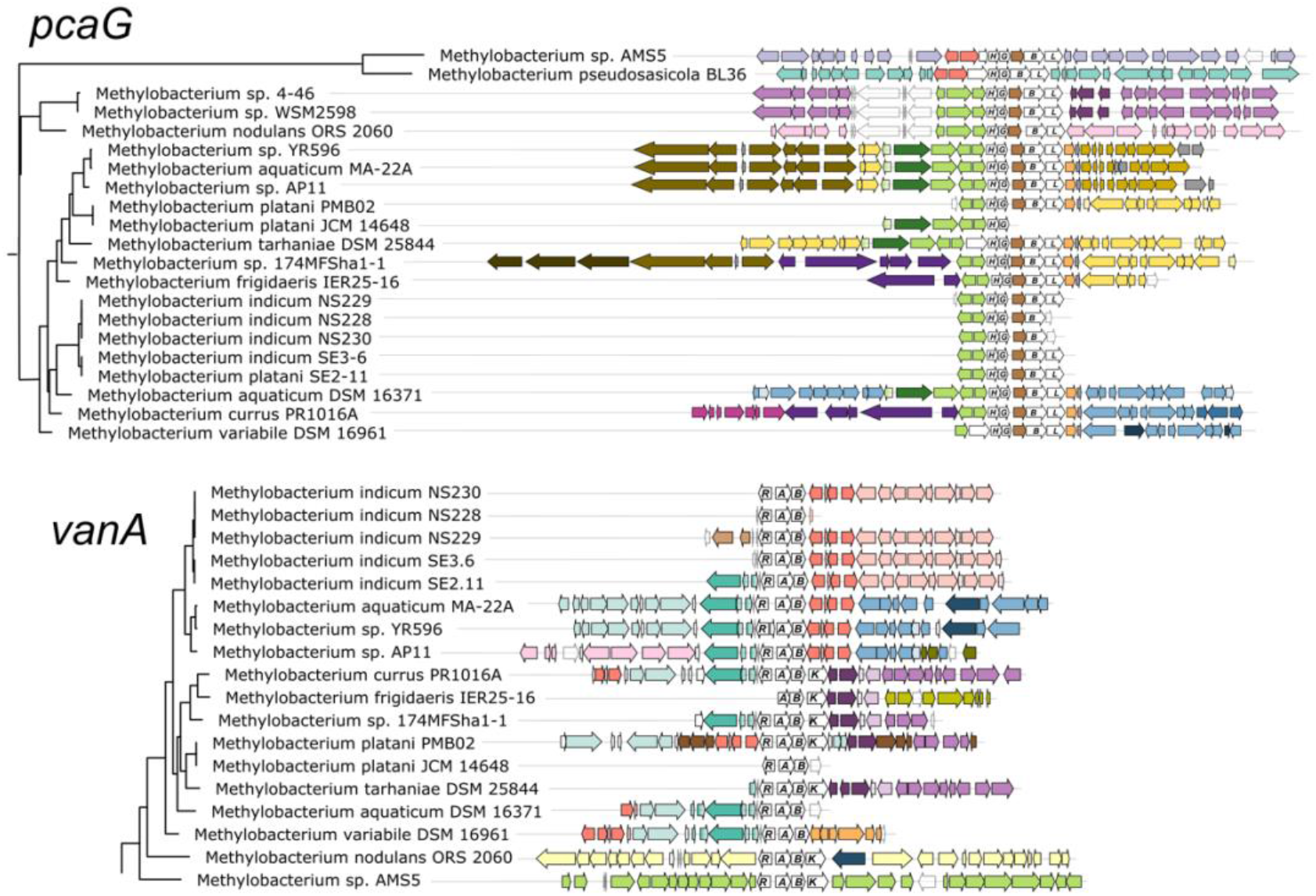
Genomic context of aromatic catabolism genes in *Methylobacterium* species supports a common evolutionary origin within the *M. aquaticum/nodulans* clades and separate origin in *M. pseudosasiacola* and *Methylobacterium* sp. AMS-5. Genomic regions (from fully assembled genomes) or scaffolds (from partially assembled genomes) showing the neighborhoods of *pcaG* and *vanA* were aligned at the genes of interest. Gene phylogenies are excerpted from Fig. 2. Genes of interest and their operons are shown in white with gene names labeled. Within each of the two alignments, homologous regions shared across multiple genomes are shown in the same color to facilitate comparison among genomes. In most cases, members of the *M. aquaticum* clade share some synteny in the regions surrounding the *van* and *pca* gene clusters, with the degree of synteny roughly paralleling their phylogenetic distance. Exceptions indicate likely cases of horizontal gene transfer, in agreement with Fig. 3.

The genomic contexts of *pcaG* and *vanA* (Fig. 3), as well as *pobA* and *ech* (Supplementary Figure 4), exhibited patterns that agreed overall with the phylogenies described above. Closely related strains shared common genes in the regions near the genes of interest, with differences among the strains increasing with increasing distance from the genes of interest. Also consistent with phylogeny, *Methylobacterium* sp. AMS5 and *M. pseudosasiacola* shared no commonalities with members of clade C or with each other in the neighborhoods surrounding any of the genes of interest. In addition, these were the only two organisms in which *pobA* and *pobR* were located adjacent to the *pca* gene cluster; this contrasts with their arrangement in the other *Methylobacterium* and in most previously described organisms, where the *pca* genes are not co-located with any of the other genes of interest in this study (Supplementary Tables 1, 2). However, PCA, the reaction product of the enzyme encoded by *pobA*, is the substrate for the gene encoded by *pcaGH*; the proximity of the genes therefore suggests that if these two organisms gained the *pca* genes via horizontal transfer, the *pob* genes may have been transferred simultaneously as part of a catabolic island. We found further support for the catabolic island hypothesis in the proximity of *vanA* and *ech* to the *pca* genes in *Methylobacterium* sp. AMS5.

### 3.4 *Methylobacterium* sp. AMS5 carries a catabolic island conferring the ability to degrade lignin-derived aromatic compounds, in an *M. extorquens*-like genome

The co-localization of all four aromatic catabolism genes of interest in the genome of *Methylobacterium* sp. AMS5 prompted us to examine the region more closely. AMS5 was originally isolated in 2011 from the stem of a hypernodulating strain of soybean and, like other organisms in the group not formally described, is currently named as a *Methylobacterium* (Anda et al., 2011), but is very closely related to the model organism *M. extorquens* PA1 and therefore likely to belong to the *Methylorubrum* group (Fig. 1). We aligned the two genomes, as well as that of another clade member, *M. zatmanii* PSBB041, at the region where the aromatic catabolism genes are located (Fig. 4). The comparison revealed that the region appears to be prone to the insertion and excision of mobile genetic elements.

**Figure 4.**
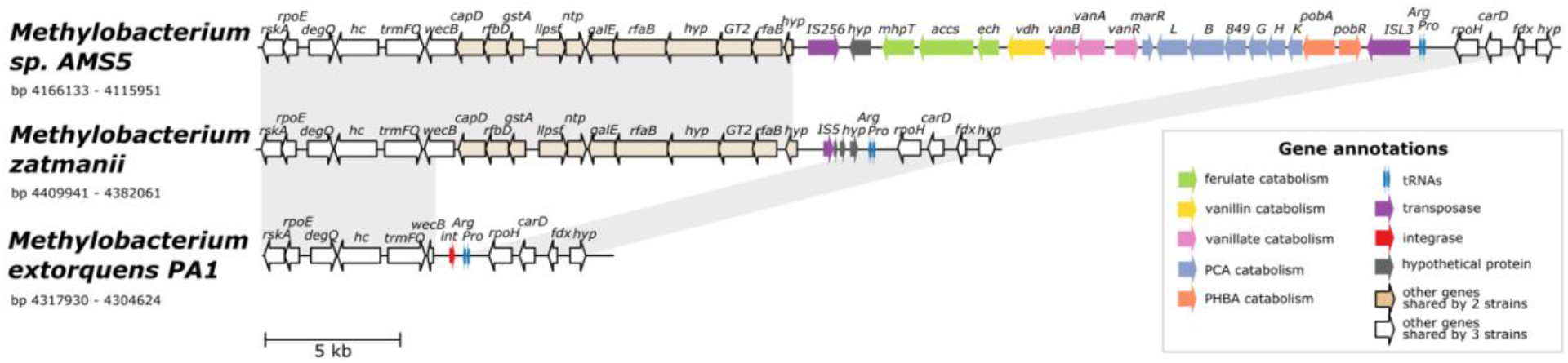
Alignment of three genomes from the *extorquens* clade reveals that *Methylobacterium* sp. AMS-5 harbors a catabolic island conferring the ability to degrade several aromatic substrates, in a genomic region prone to insertions and deletions. The genome of *Methylobacterium* sp. AMS5 is shown aligned with that of two close relatives, *M. zatmanii* PSBB041 and *M. extorquens* PA1. Genes are color-coded to indicate either function or commonality between genomes, as indicated by the key. Light gray shading between pairs of species connects regions shared by both. See Table S3 for full annotations corresponding to the short gene codes.

At the locus where *M. zatmanii* carries a gene encoding a putative IS5-type transposase (possibly nonfunctional due to a frameshift with an internal stop codon), AMS5 contained an additional 22-kb region: a catabolic island containing genes that appear to confer the full ability to degrade ferulate, vanillate, vanillin, *p*-hydroxybenzoate, and protocatechuate, flanked by two different transposase genes from the MULE superfamily. On the 3’ side of this region, the two strains share 11 genes (14 kb), many related to nucleotide sugar metabolism, that are also not present in PA1 (Fig. 4), or in any of the other *Methylobacterium* species surveyed in this study. The corresponding locus in the PA1 genome carries only a putative site-specific integrase/recombinase, and this gene is present only in members of the *extorquens* clade (clade B). The gene adjacent to the integrase encodes UDP-*N*-acetylglucosamine 2-epimerase (*wecB*); the sequence appears to be complete in *M. zatmanii* (1,137bp) but truncated in PA1 (missing 912 bp from the 3’ end). In all three genomes, the region of variability lies immediately at the 3’ end of the genes for tRNA-Pro and tRNA-Arg. Due to the extensive genome rearrangement that is common within the *Methylobacterium* (Vuilleumier et al., 2009; Lee and Marx, 2012), we were unable to identify a syntenic region in species of clades A or C for comparison.

The presence of the truncated *wecB* gene in PA1 suggests that the region of sugar metabolism genes may originally have been present but were excised, whereas the arrangement of genes in the catabolic island suggest that the island was acquired into a *M. zatmanii*-like genomic background. The origins of the catabolic island itself are uncertain; as shown above (Fig. 2), there appear to be diverse phylogenetic origins represented among the different genes within the pathway, and BLAST search for the entire region in the NCBI Nucleotide database found no other organisms outside of the *Methylobacterium* containing all genes in the same order. We could detect no genes in the region relating to replication or conjugation as might be expected in an integrative and conjugative element (ICE) (Wozniak and Waldor, 2010). Furthermore, no significant difference in tetranucleotide composition or in GC content was found between the inserted region (GC% = 71.4) and the full genome (GC% = 71.1) (Supplementary Figure 5), indicating that the catabolic island was likely either transferred from an unknown closely-related organism, or has been present in the genome long enough for amelioration (Lawrence and Ochman, 1997).

### 3.5 Genome content predicts ability to grow on aromatic compounds in most *Methylobacterium* strains

We next wanted to assess whether *Methylobacterium* strains carrying genes for the catabolism of methoxylated aromatic compounds could indeed use those compounds as growth substrates. We chose 8 species from the *M. aquaticum/nodulans* clades, shown in Figure 5 (and Supplementary Table 1). All of these strains carry all four genes of interest except *Methylobacterium* sp. 4-46, which lacks *vanA* and so was predicted not to grow on vanillate or ferulate (Fig. 1). We conducted a series of growth experiments on defined mineral medium with substrates of each of the four gene products of interest, as well as several other substrates for context: methanol (a simple C1 substrate), succinate (a known growth substrate for all the strains tested), glucose (used by only some *Methylobacterium* species), benzoic acid (another aromatic acid with a degradation pathway separate from the beta-ketoadipate pathway (Moreno and Rojo, 2008)), and Kraft lignin (a soluble form of polymeric lignin). All strains grew on succinate and none on lignin; growth on PHBA, PCA, vanillate, and ferulate was as expected for all strains given their genome content, with the exceptions that *M. variabile* did not grow on vanillate and *M. nodulans* did not grow on ferulate (Fig. 5; Supplementary Tables 3, 4). Because a number of factors may lead to lack of growth, further tests in different culture conditions would be necessary to further explore these strains’ ability to grow on these substrates.

**Figure 5.**
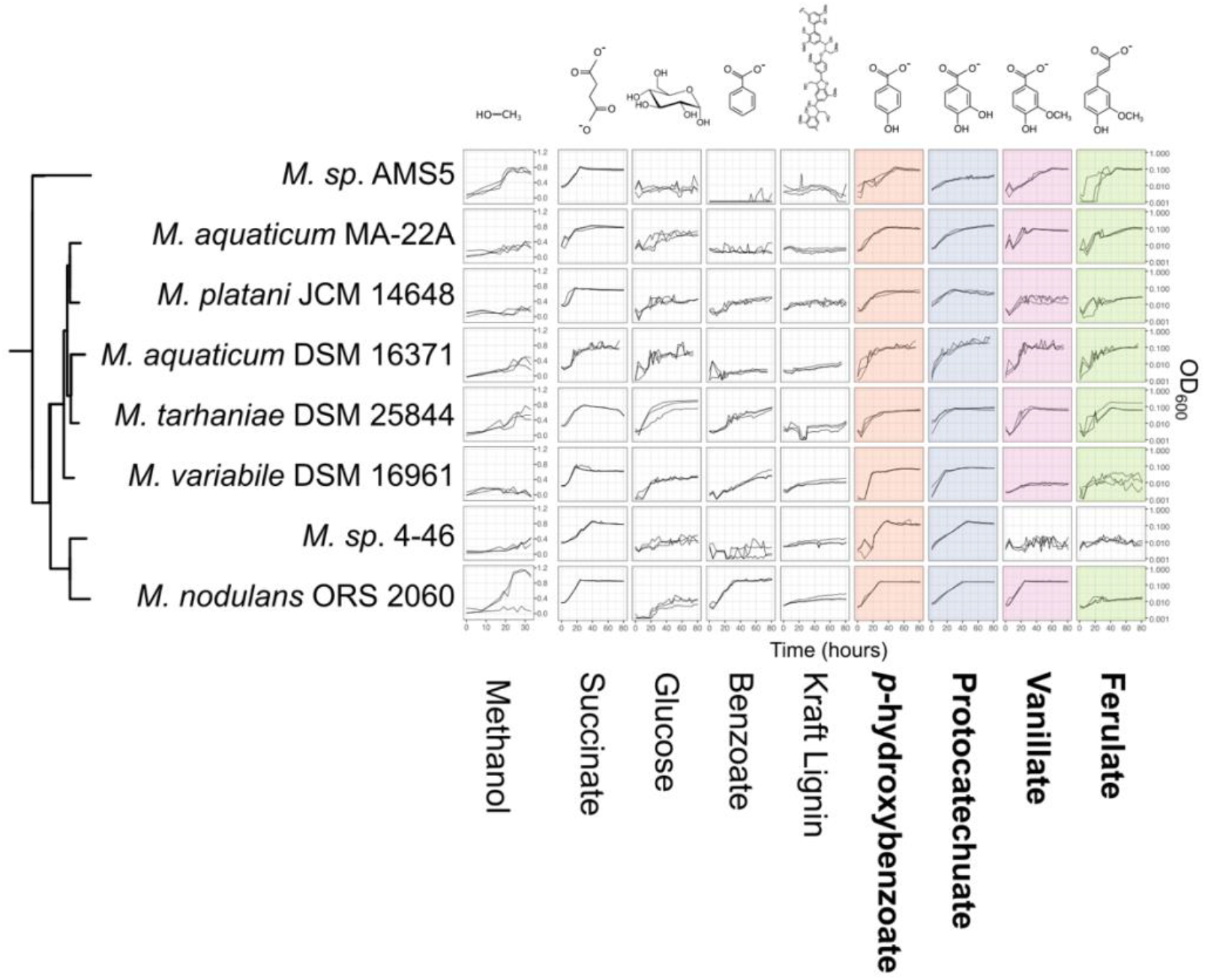
Growth of *Methylobacterium* species on aromatic compounds is predicted by genome content in most cases. *Methylobacterium* species were grown in defined mineral medium with a single carbon substrate; all conditions contained 16 mM of carbon. Substrates corresponding to the four degradation pathways featured in this study are outlined labeled in bold type, and presence of the corresponding genes in each strain’s genome is indicated by colored shading (as in Fig. 1). Lines show growth curves from three biological replicates. Note that the y-axis for methanol is on a different scale, as these cultures were grown in different vessels. There were only 3 cases (out of 32) in which organisms had the genes associated with degradation of a given substrate but showed little or no growth. There were no cases in which organisms grew on substrates for which they did not have the associated genes. Source data are available in Supplementary Tables 4 and 5.

### 3.6 Non-methylotrophs excrete formaldehyde during vanillate growth, whereas *Methylobacterium* do not

We were especially interested to ask whether methylotrophs are different from non-methylotrophs in their ability to cope with the formaldehyde produced during vanillate metabolism. As mentioned above, it has been recognized by multiple researchers that the formaldehyde released by vanillate monooxygenase is a burden to organisms growing on vanillate, and we have previously observed formaldehyde accumulation during vanillate utilization by *Pseudomonas putida* KT2440 (Lee et al., 2021). We therefore measured formaldehyde in the medium during vanillate metabolism by *P. putida* and *Rhodoccus jostii* RHA1, two well-studied lignin-degrading bacteria that, unlike *Methylobacterium*, do not utilize C1 compouds for growth. We assayed each strain on vanillate as well as on PCA as a control compound, as PCA is the direct product of vanillate demethylation and therefore involves the same metabolism except for the effect of formaldehyde. In both strains, when growth occurred on vanillate, formaldehyde accumulated in the medium concomitant with the increase in optical density, peaking at concentrations of 0.94 mM (Fig. 6A, Supplementary Table 5). When cultures entered stationary phase (approximately 10 hours for *P. putida*, 15 hours for *R. jostii*), formaldehyde concentrations began to decrease, ultimately returning to below the detection limit. No formaldehyde was detected at any time during growth on PCA. Moreover, growth on PCA was faster: stationary phase was reached at 7 hours for *P. putida* and 10 hours for *R. jostii*. These results suggest that while both *P. putida* and *R. jostii* can oxidize formaldehyde, removal is slower than production.

**Figure 6.**
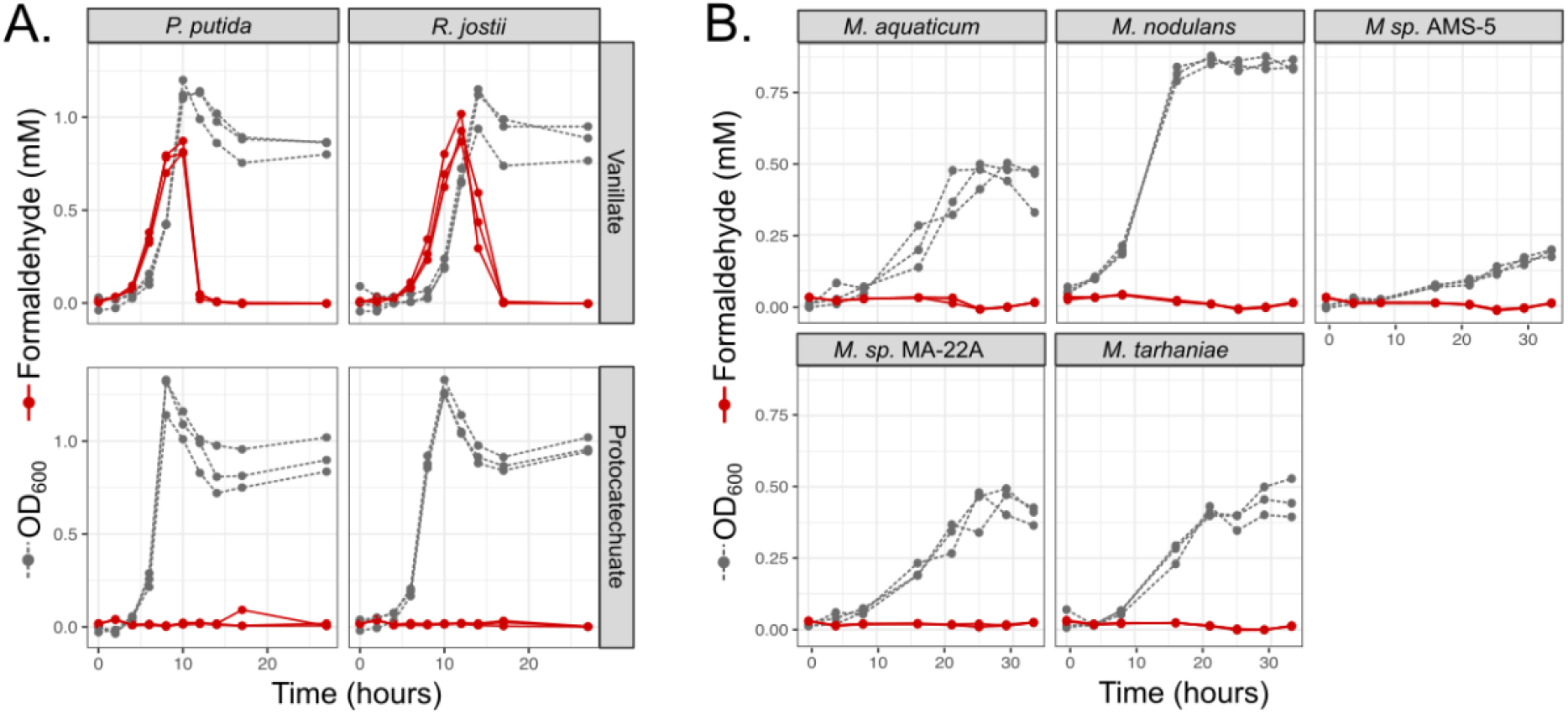
Non-methylotrophic lignin degraders *Pseudomonas putida* and *Rhodococcus jostii* produce formaldehyde when growing on methoxy-substituted aromatic compounds such as vanillate; *Methylobacterium* do not. All organisms were grown in defined mineral medium with vanillate or protocatechuate (PCA) as a sole carbon source; growth was assayed by optical density at 600 nm (gray symbols and dashed lines) and formaldehyde in the medium was measured by a colorimetric assay (red symbols and solid lines). Each line represents one biological replicate. (A) *P. putida* and *R. jostii* accumulate formaldehyde transiently in the medium when growing on vanillate, but not on the non-methoxylated compound PCA, and growth is slower on vanillate. (B) All *Methylobacterium* species tested grew on vanillate without producing measurable formaldehyde. Source data are available in Supplementary Table 6.

We conducted a similar experiment on the five strains of *Methylobacterium* showing the greatest growth on vanillate in our laboratory conditions: *M. aquaticum*, *M. nodulans*, *M. tarhaniae*, *Methylobacterium* sp. AMS5, and *M. aquaticum* MA-22A. We incubated all cultures with vanillate for 33 hours, long enough for AMS5 to show marked growth and all other species to consume all the substrate and reach stationary phase. No detectable formaldehyde was produced at any time (Fig. 6B, Supplementary Table 5). These results suggest that *Methylobacterium* species are able to consume the formaldehyde generated from vanillate demethylation as rapidly as it is produced, likely via the same pathways by which the formaldehyde generated from methanol is used for energy generation and biosynthesis (Marx et al., 2005; Crowther et al., 2008).

### 3.7 *Methylobacterium*-derived *vanA* reads in published metagenomes are predominantly from the *M. nodulans* cluster, and most numerous in soil

Aside from the environments in which they were isolated, there exists scant information on the ecological niches of the aromatic-degrading *Methylobacterium* clades—for instance, whether they are more likely than the other *Methylobacterium* to inhabit ecosystems rich in lignin, such as soil or the rhizosphere. We therefore sought to learn more about the prevalence and abundance of *vanA*-carrying *Methylobacterium* species in the environment by querying the publicly available metagenome datasets on the JGI IMG/M database. The distinctiveness of the *Methylobacterium* genes within the phylogeny of known *vanA* sequences (Fig. 2) makes it possible to deduce phylogeny from DNA sequence and *vanA* is present as a single copy in most genomes in which it is found.

Our study set comprised 1,651 metagenomes, with a total assembled gene count of 5.60×10^9^; we retrieved 317,816 scaffolds carrying *vanA*, of which 348 had *Methylobacterium* or *Methylorubrum* as their IMG Phylogenetic Lineage Assignment (a frequency of 0.11%) (Supplementary Tables 6, 7, 8). We then used pplacer (Matsen et al., 2010) (RRID:SCR_004737) for phylogenetic placement of those reads onto a reference tree of *vanA* genes from known *Methylobacterium*, and found that 182 of them actually clustered with the *Methylobacterium*; the others fell outside the *Methylobacterium* and were therefore omitted from further study (Supplementary Figure 1). Of those 182 *Methylobacterium vanA*, 114 (63%) clustered with the *M. nodulans vanA* gene; 31 genes (17%) with *Methylobacterium* sp. AMS5, and 37 genes (20%) fell into the *M. aquaticum*-like clade (Fig. 7). Overall, they were most commonly found in samples associated with soil (43%) and roots (40%); only 10% were found in leaf samples.

**Figure 7.**
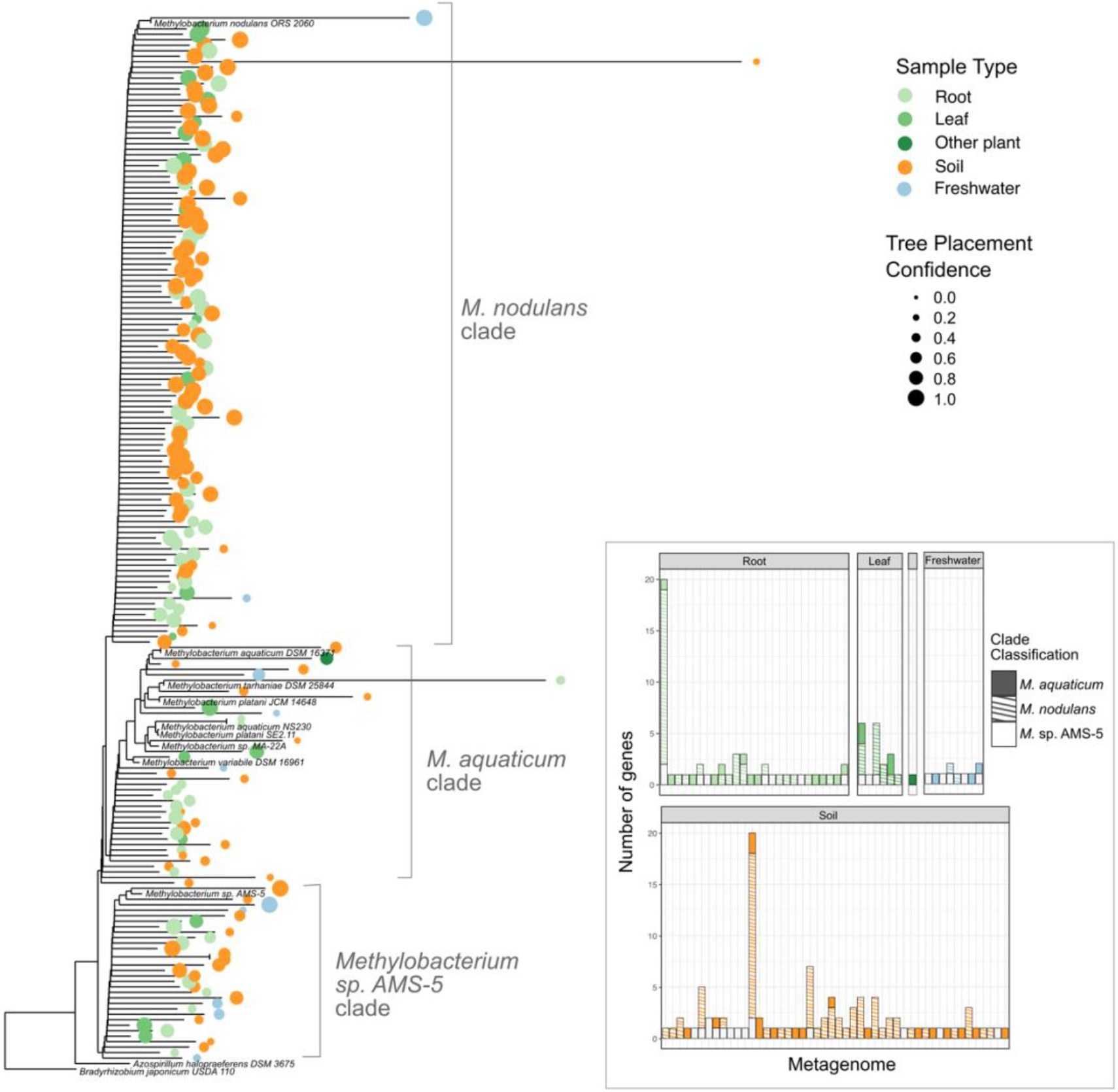
Among *Methylobacterium vanA* sequences in published metagenomes, *M. nodulans*-like sequences are the most abundant. Sequences of 182 *Methylobacterium*-associated *vanA* gene fragments found in published metagenomes were added using pplacer to a phylogeny of full-length *vanA* genes sequenced from reference genomes. 166 genes that were classified by IMG as *Methylobacterium* but clustered outside of the genus are not shown. Colored dots indicate the ecosystem type from which the metagenome sample originated, with size scaled to the likelihood weight ratio of the pplacer classification as a measure of confidence in the placement. Inset bar plot: abundance of *Methylobacterium vanA* genes, shaded by clade and colored by sample type, in each metagenome in which they were found. Each bar represents one metagenome (genome IDs not shown).

### 3.8 *Methylobacterium* metagenome reads show ecological differences among clades

To place these findings in context, we carried out a survey for *Methylobacterium* metagenome reads across all ecosystem types. We used the Phylogenetic Distribution of Genes in Metagenome tool (Markowitz et al., 2012; JGI IMG, 2017; Chen et al., 2019) to query all published metagenomes for reads with 90% BLAST similarity to each of 18 genomes from the *Methylobacteriaceae*. We observed a pattern in the samples that had BLAST hits to the query genomes suggesting ecological differences among the clades (Fig. 8; Supplementary Figure 6; Supplementary Tables 9, 10, 11).

**Figure 8.**
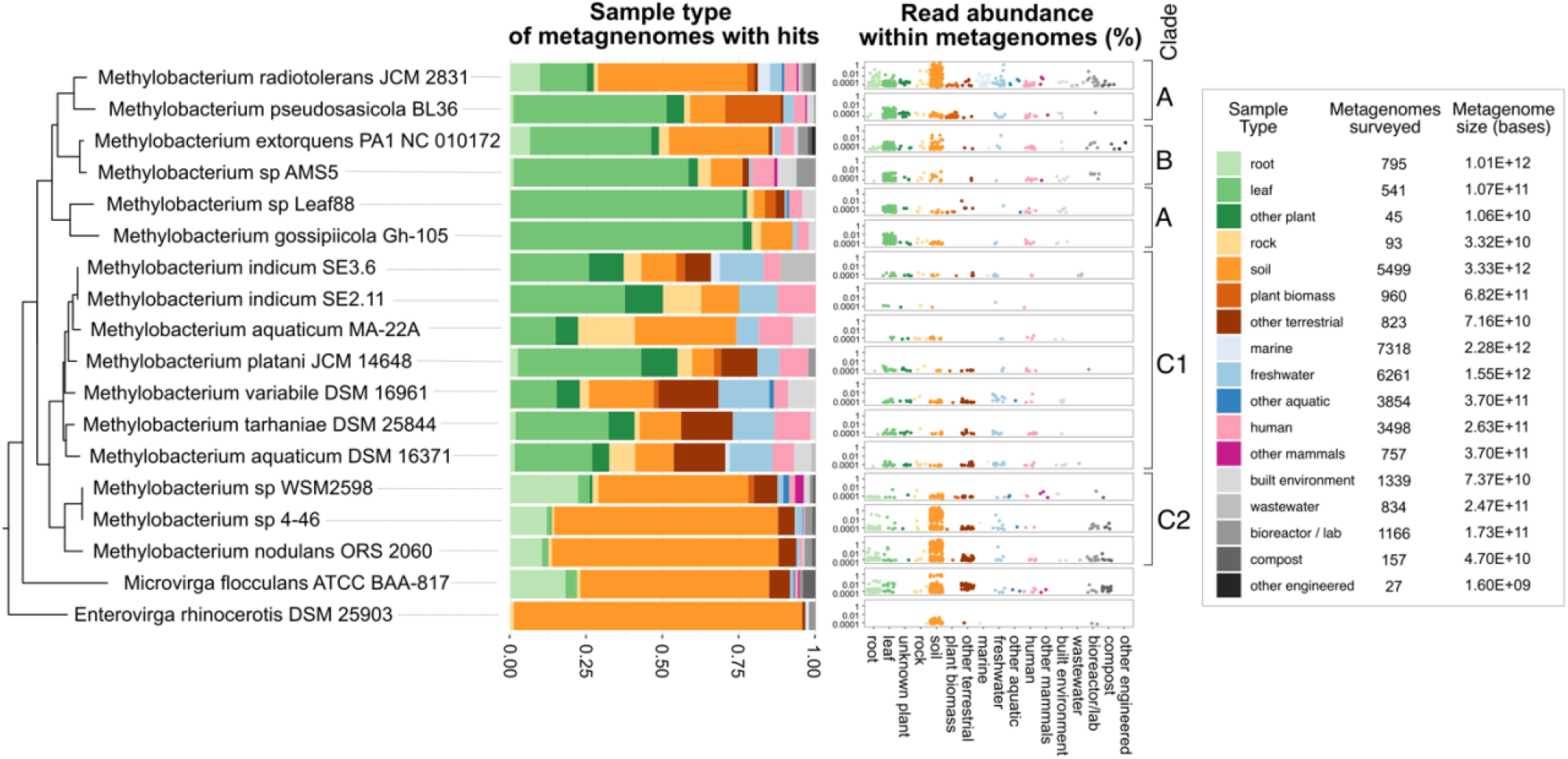
*Methylobacterium* strains vary substantially in their distribution among ecosystems; not all strains are phyllosphere-associated. Published reads from 33,967 metagenomes were surveyed for BLAST hits with 90% or greater similarity to the genomes of 18 strains from the *Methylobacteriaceae*, using the Phylogenetic Distribution of Genes in Metagenome tool in IMG. Each metagenome was classified into one of 17 ecological sample types. For each strain, the barplot shows the distribution of BLAST hits among those sample types. In the dotplots, each dot represents one metagenome, with height indicating the abundance of reads matching the strain’s genome within that metagenome. Points are color-coded, and clustered along the x-axis, by sample type. For context, the number of metagenomes, and total number of bases surveyed (in all metagenomes) is provided for each sample type. Phylogeny of the *rpoB* gene of the target species (as in Supplementary Figure 2) is shown at left.

For example, the hits to query genomes from clades A and B were generally found with higher frequency in metagenomes from leaf samples than from soil (for *Methylobacterium sp*. Leaf88 and *M. gossipiicola*, respectively, 4% and 10% of hits were from the soil, whereas 76% and 76% of were from leaf samples). In contrast, hits to query genomes from clade C2 were found primarily in soil and rarely in leaf samples (for *M. nodulans*, *Methylobacterium sp*. 4-46, and *Methylobacterium sp*. WSM2598, respectively, 74%, 73%, and 49% of hits were from the soil, whereas 2%, 1%, and 4% were from leaf samples). Clade C1 differed from the others in that there were fewer hits to these query genomes (between 8 and 71 hits per genome, in contrast to 96-487 hits for clades A-B and 104-523 hits for clade C2), and those hits more evenly distributed across all the sample types surveyed. These results point to ecological differences between the two clades of *Methylobacterium* capable of aromatic catabolism (clades C1 and C2). However, in general they agree with the survey of *Methylobacterium vanA* metagenome reads, suggesting that these two clades are more likely to be found in soil and root environments, and less likely to be found in the phyllosphere, than other clades of *Methylobacterium*.

## 4 Discussion

We have provided the first description of the methoxydotrophic growth of *Methylobacterium* on lignin-derived methoxylated aromatic acids. *Methylobacterium* species with this capability can use vanillate as a sole carbon substrate, without the transient formaldehyde accumulation in the environment observed in vanillate degraders that do not grow on C1 compounds. The genes encoding aromatic catabolism are found in almost all genome-sequenced members of clades C1 and C2 (those related to *M. aquaticum* and *M. nodulans*); they were largely absent in clades A and B (the *M. radiotolerans* and *M. extorquens* clades), but acquisition by horizontal gene transfer may have taken place in some species of those clades.

These findings shed new light on the ecology and evolution of the *Methylobacterium. Methylobacterium* have traditionally been of interest due to their ability to use single-carbon compounds as a sole source of carbon and energy; indeed, as their name suggests, methylotrophy has been one of the criteria used for classification. However, most work thus far has focused on methanol, methylamine, or formate as growth substrates; to our knowledge, ours is the first study to address whether methoxylated aromatic compounds might serve as a source for those single-carbon compounds. In fact, very few studies have addressed the possibility of *Methylobacterium* species using aromatic compounds as growth substrates at all.

As organisms known for their association with plants, it is possible that *Methylobacterium* play a role in the bacterial lignin-degrading community. In the environment, these organisms may depend upon aromatic acids released by the action of other lignin-degrading organisms, which have been found to be a prominent component of leaf litter leachate (Kuiters, 1990). Alternatively, *Methylobacterium* may encounter methoxylated aromatic acids primarily in plant root exudates (Zhalnina et al., 2018). In addition to acting as a growth substrate, aromatic acids play an important role in plant-microbe signaling: vanillate, ferulate, *p*-hydroxybenzoate, and protocatechuate all influence on the process and productivity of root nodulation by other members of the *Rhizobiales* (Seneviratne and Jayasinghearachchi, 2003; Mandal et al., 2010). This is significant in light of the fact that the members of the *M. nodulans* clade were, as the species name suggests, isolated from root and stem nodules of their plant hosts (Fleischman and Kramer, 1998; Jourand, 2004); and that we found this clade to be the most abundant among the *Methylobacterium vanA* genes found in the environment. Furthermore, *Methylobacterium* sp. AMS5 was isolated in a study on soybean epiphytes that are particularly responsive to host nodulation phenotype (Anda et al., 2011). It is likely that at least part of the importance of aromatic catabolism in *Methylobacterium* is to facilitate the relationships of these plant-associated organisms with their hosts. A link between aromatic catabolism and plant-microbe symbioses could help to explain our finding that *vanA* sequences related to those from *M. nodulans* and *Methylobacterium* sp. AMS5 are dominant among the *Methylobacterium vanA* sequences found in the environment despite the fact that there are few genome-sequenced representatives from those groups.

Perhaps the most compelling results from this study are the new insights into the evolution of the *Methylobacterium* group. One element is the absence of the *vanA* gene in *Methylobacterium* sp. 4-46 and sp. WSM298, the only two species we found in clade C to lack the capacity for methoxydotrophic growth. These species do carry genes for transforming ferulate to vanillin (*ech*) and vanillin to vanillate (*vdh*), which may be remnants from predecessors that were able to metabolize vanillate but lost the capability. Notably, *Methylobacterium* sp. 4-46 is also one of very few *Methylobacterium* species reported not to be capable of methylotrophic growth (Green and Ardley, 2018). Similarly to other strains described in (Alessa et al., 2021), we have found that *Methylobacterium* sp. 4-46 is indeed capable of growth on methanol, but only in the presence of LaCl_3_ (Supplementary Figure 7, Supplementary Table 12). This suggests the involvement of a XoxF-type methanol dehydrogenase (Skovran and Martinez-Gomez, 2015), and possibly a different role for methanol oxidation in this organism’s ecology. Given our hypothesis that methanol oxidation and vanillate demethylation require the same pathways for metabolizing the formaldehyde produced, it is possible that the loss of *vanA* in *Methylobacterium* sp. 4-46 might be related to its different style of methylotrophy. We have also observed that this and several non-*Methylobacterium* lignin-degrading species possess the genes to use PCA (the aromatic product of vanillate demethylation) but not the methoxy group (Fig. 1). Yet we have found no species in which the reverse is true, although it would theoretically be possible for a methylotroph with *van* genes but no *pca* genes to carry out methoxydotrophic growth by utilizing only the methoxy group of vanillate.

The other unexpected evolutionary finding relates to the acquisition of aromatic catabolism genes by horizontal transfer in two *Methylobacterium* species from outside of clade C. The discovery of a catabolic island in *Methylobacterium* sp. AMS5 is itself not unusual; *Methylobacterium* and *Methylorubrum* species have long been recognized to carry an abundance of IS (insertion sequence) elements, and it has been postulated that the associated genome rearrangements and horizontal gene transfer associated are important mechanisms of evolution in the group (Schmid-Appert et al., 1997; Vuilleumier et al., 2009; Nayak and Marx, 2015). Relevant to the present study is the prior observation that diverse features of the genomic background—and not necessarily those predicted by phylogeny—influence whether a newly introduced set of genes is immediately useful to the recipient organism (Michener et al., 2014, 2016). Are there particular features of the *M. pseudosasiacola* and *Methylobacterium* sp. AMS5 genomes that allowed them to acquire the capacity for the degradation of methoxylated aromatics when no other known members of their clades did, or is the maintenance of this genomic capability the result of selective pressure specific to their ecological niche? Further work on the ecology of AMS5 and the biology of aromatic catabolism in *Methylobacterium* is necessary to address these questions.

This study has benefited from the wealth of knowledge that already exists on pathways for the degradation of lignin-derived aromatics in other microorganisms, and on methylotrophic metabolism in *Methylobacterium*, to deduce the likely fate of the methoxy group from vanillate. We found that in most cases, the gene annotations in IMG enabled us to correctly predict the substrates each strain could grow on (the exceptions were cases of no growth, and we cannot rule out the possibility that growth could occur under different conditions). However, we did find some novel features of the *Methylobacterium* pathways: almost all the strains we studied appear have no homologs of *pcaI, pcaJ, or pcaF* encoded within the *pca* gene cluster. It is possible that the functions of these genes are carried out by homologs located elsewhere in the genome, as has been found in some other organisms (Parke, 1997). A second possibility is raised by a previous study that carried out enzymatic screening and active site modeling on the DUF849 family of genes (Bastard et al., 2014): several of the DUF849 genes found in these *Methylobacterium* gene clusters were classified as beta-keto acid cleavage enzymes (G4 BKACE) predicted to act on betaketoadipate, raising the possibility that they might carry out the *pcaI/pcaJ* function and thus constitute a novel variant of the already diverse family of beta-ketoadipate pathway configurations (Parke, 1997; Buchan et al., 2004).

Our survey of where methoxydotrophic vanillate-utilizing *Methylobacterium* might be found in the environment suggests that they are more prominent in soil than in leaf environments. This is a notable given that reports on the *Methylobacterium* often focus on the phyllosphere as a characteristic habitat (e.g., (Iguchi et al., 2015; Alessa et al., 2021)). Our findings relating to *Methylobacterium*-associated *vanA* genes and *Methylobacterium* reads in metagenomes are subject to numerous caveats, including variability in extraction and sequencing methods and metadata annotation in public datasets, and the limitations in our ability to deduce taxonomy from short sequencing reads. Moreover, the methods we used are unable to capture the role of the horizontal gene transfer that is likely important for conferring aromatic catabolism to organisms such as *Methylobacterium* sp. AMS5. To gain true insight into the ecology of aromatic catabolism in this group will require more targeted sequencing efforts, as well as isolation and cultivation. However, this work provides a solid motivation for future studies, by demonstrating that *Methylobacterium* have diverse ecological patterns across their phylogeny, and that part of that diversity may be a soil-associated niche for aromatic-degrading strains.

## Supporting information

Supplementary Datasheets

## 5 Data Availability Statement

The datasets generated for this study are provided in the Supplementary Material. The publicly available datasets analyzed for this study can be found in the Integrated Microbial Genomes and Microbiomes database (IMG/M) (Chen et al., 2019); accession numbers of all metagenomes, scaffolds, and genes used are provided in the Supplementary Material.

## 6 Author Contributions

JAL: Conceptualization, Methodology, Investigation, Data Curation, Writing - Original Draft, Visualization. SS: Methodology, Resources, Writing - Review & Editing. CUM: Methodology, Resources, Writing - Review & Editing, Supervision, Funding acquisition.

## 9 Acknowledgments

We are grateful to Nicholas Shevalier, Alyssa Baugh, and Tomislav Ticak for their assistance with experiments and support during the scientific process. We thank Armando MacDonald for technical assistance with measurements, Akio Tani for providing the *M. aquaticum* MA-22A culture, Andrea Lubbe and Trent Northen for the *P. putida* and *R. jostii* cultures, and José de la Torre for assistance with calculating genome signatures. We are thankful to Dipti Nayak, Mete Yuksel, Elizabeth Winters, Brittany Baker, and Tomislav Ticak for their valuable feedback on the manuscript.

## 10 Supplementary Material

### 10.1 Supplementary Figures

1. IMG phylogenetic lineage annotations do not match pplacer phylogenetic placement of *vanA* genes.

2. Phylogeny of all *Methylobacterium* and *Methylorubrum* strains included in this study, based on the *rpoB* gene.

3. Phylogeny of vanillate monooxygenase (*vanA*) genes with other COG 4638 (Phenylpropionate dioxygenase or related ring-hydroxylating dioxygenase, large terminal subunit) genes from *M. nodulans, M. indicum* NS230, *M. aquaticum* DSM 16371, *M. pseudosasiacola*, and *M. extorquens* PA1.

4. Genomic context of aromatic catabolism genes *pobA* and *ech* in *Methylobacterium* species.

5. *Methylobacterium sp*. AMS5 catabolic island has similar GC content and tetranucleotide composition to the rest of the genome.

6. Ecological distribution of metagenomes surveyed for *Methylobacterium* reads, for comparison with Fig. 8.

7. *Methylobacterium* sp. 4-46 grows on methanol only when lanthanum is provided.

### 10.2 Supplementary Tables

1: *Methylobacterium* strains included in this study, and locus tags of genes analyzed [xlsx]

2: Reference organisms included in Figures 1 and 2, and locus tags of genes analyzed [xlsx]

3: Growth data for *Methylobacterium* strains on diverse compounds, shown in Fig. 5 [csv] Time = incubation time in hours. Species = abbreviated strain name. Substrate = carbon substrate. Well = location of the replicate in a multiwell culture plate. OD = optical density at 600 nm.

4: Growth data for *Methylobacterium* strains on methanol, shown in Fig. 5 [csv] Time = incubation time in hours. Species = abbreviated strain name. Substrate = carbon substrate. Rep = replicate designation (A, B, or C). OD = optical density at 600 nm.

5: Full OD and formaldehyde data from growth experiments on PCA and vanillate shown in Fig. 6 [xlsx]

6: Summary of metagenomes surveyed for *Methylobacterium vanA* (Fig. 7) [xlsx]

7: All metagenomes in which *Methylobacterium vanA* genes (shown in Fig. 7) were found [xlsx]

8: All metagenome scaffolds containing *vanA* genes and identified as *Methylobacterium*, shown in Fig. 7 [xlsx]

9: Metagenomes surveyed for BLAST hits to each of 18 *Methylobacteriaceae* query genomes [xlsx]

10: Metagenomes in which BLAST hits with 90% similarity to a *Methylobacteriaceae* query genome were found.

11: Supplementary Table 11. Summary of results from BLAST search of metagenomes for hits to each query *Methylobacteriaceae* genome, by sample type [xlsx]

12: Growth of *Methylobacterium* sp. 4-46 with and without lanthanum, shown in Supplementary Figure 7 [csv] Time = incubation time in hours. OD = optical density at 600 nm. La = with or without lanthanum (25 μM LaCl_3_). Rep = replicate designation (A, B, or C).

### 10.3 Supplementary Data Sheets

1: *atpD* gene alignment (from Fig. 1)

2: *recA* gene alignment (from Fig. 1)

3: *rpoB* gene alignment (from Fig. 1)

4: 16S rRNA gene alignment (from Fig. 1)

5: *ech* gene alignment (from Fig. 2)

6: *pcaG* gene alignment (from Fig. 2)

7: *pobA* gene alignment (from Fig. 2)

8: *vanA* gene alignment (from Fig. 2)

9: *rpoB* gene alignment, all *Methylobacterium* surveyed (from Supplementary Figure 3)

10: alignment of COG 4638 homologs (from Supplementary Figure 4)

