## Supplementary Figures for "Aerobic methoxydotrophy: growth on methoxylated aromatic compounds by *Methylobacterium*"

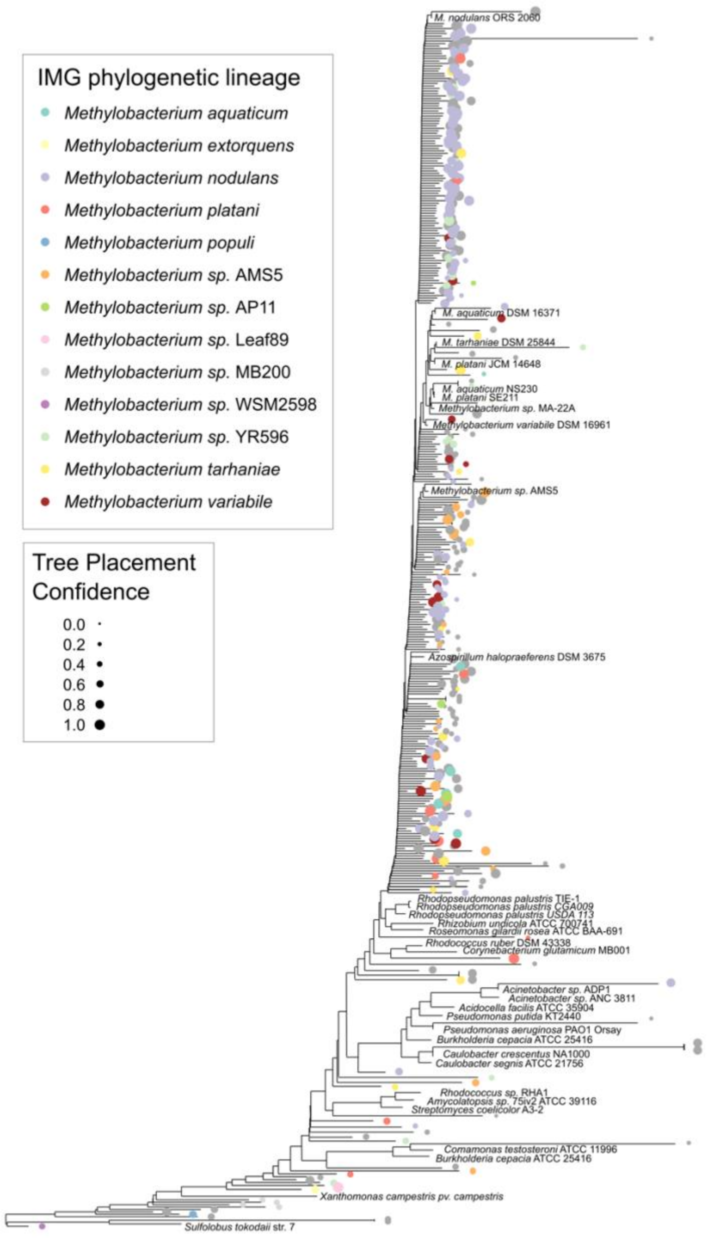


**Supplementary Figure 1**. IMG phylogenetic lineage annotations do not match pplacer phylogenetic placement of *vanA* genes. 348 of the *vanA* genes from published genomes on the IMG/M database had a phylogenetic lineage annotation of *Methylobacterium*. These genes were added to a phylogeny of *vanA* from cultured reference organisms (a subset of those from Fig. 2) using pplacer. Branches are labeled with circles color-coded according to the original IMG phylogenetic lineage; circle size is scaled to the likelihood weight ratio of the pplacer classification as a measure of confidence in the placement. Many of the gene sequences did not cluster with the *Methylobacterium*. Only those that did were analyzed for this study.

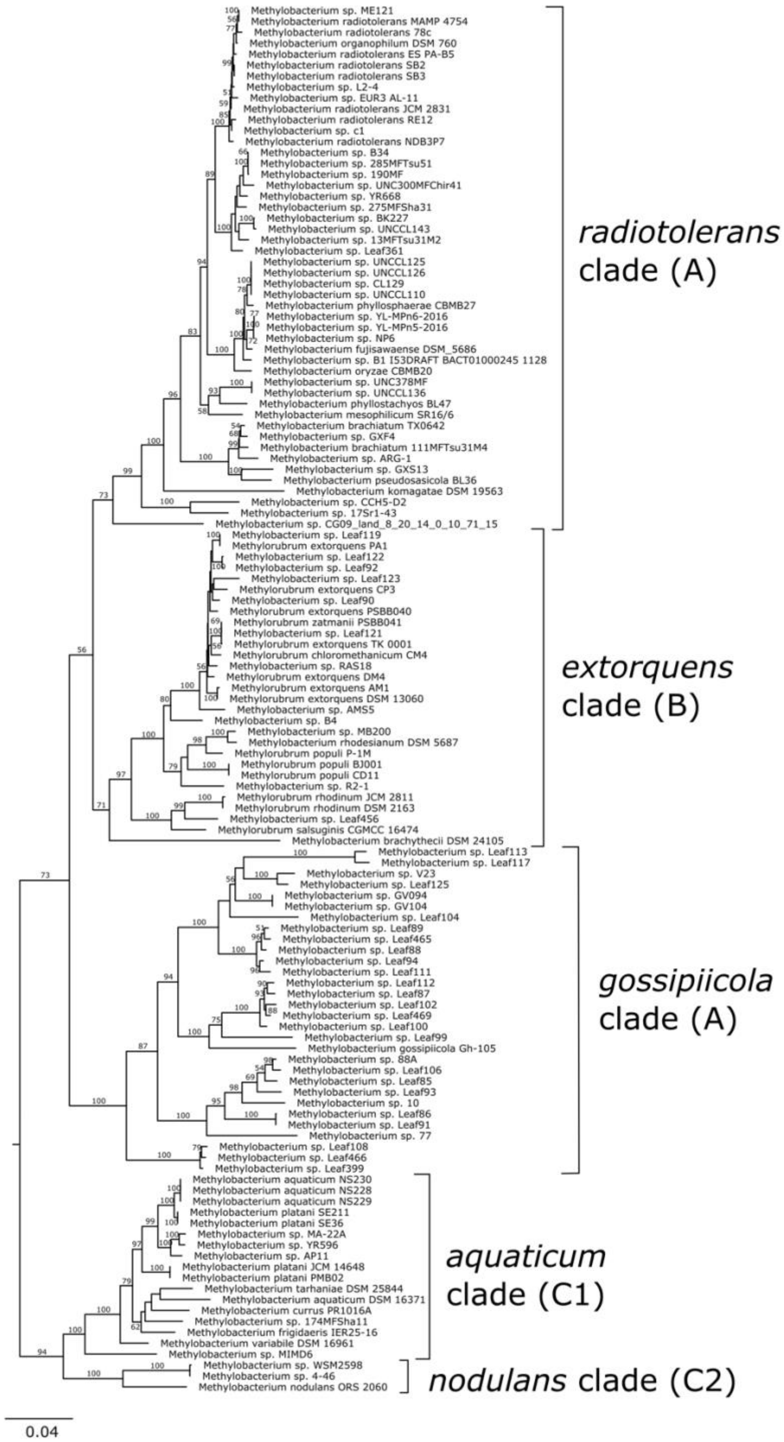


**Supplementary Figure 2.** Phylogeny of all *Methylobacterium* and *Methylorubrum* strains included in this study, based on the *rpoB* gene (encoding the beta subunit of RNA polymerase). Branch labels indicate bootstrap percentage. Clade A/B/C designations are in accordance with Green and Ardley (2018); Alessa et al. (2021); and Leducq et al. (2021). This tree is provided for context, as not all strains analyzed are displayed in Figure 1. Aligned sequence data are given in Supplementary Data Sheet 9.

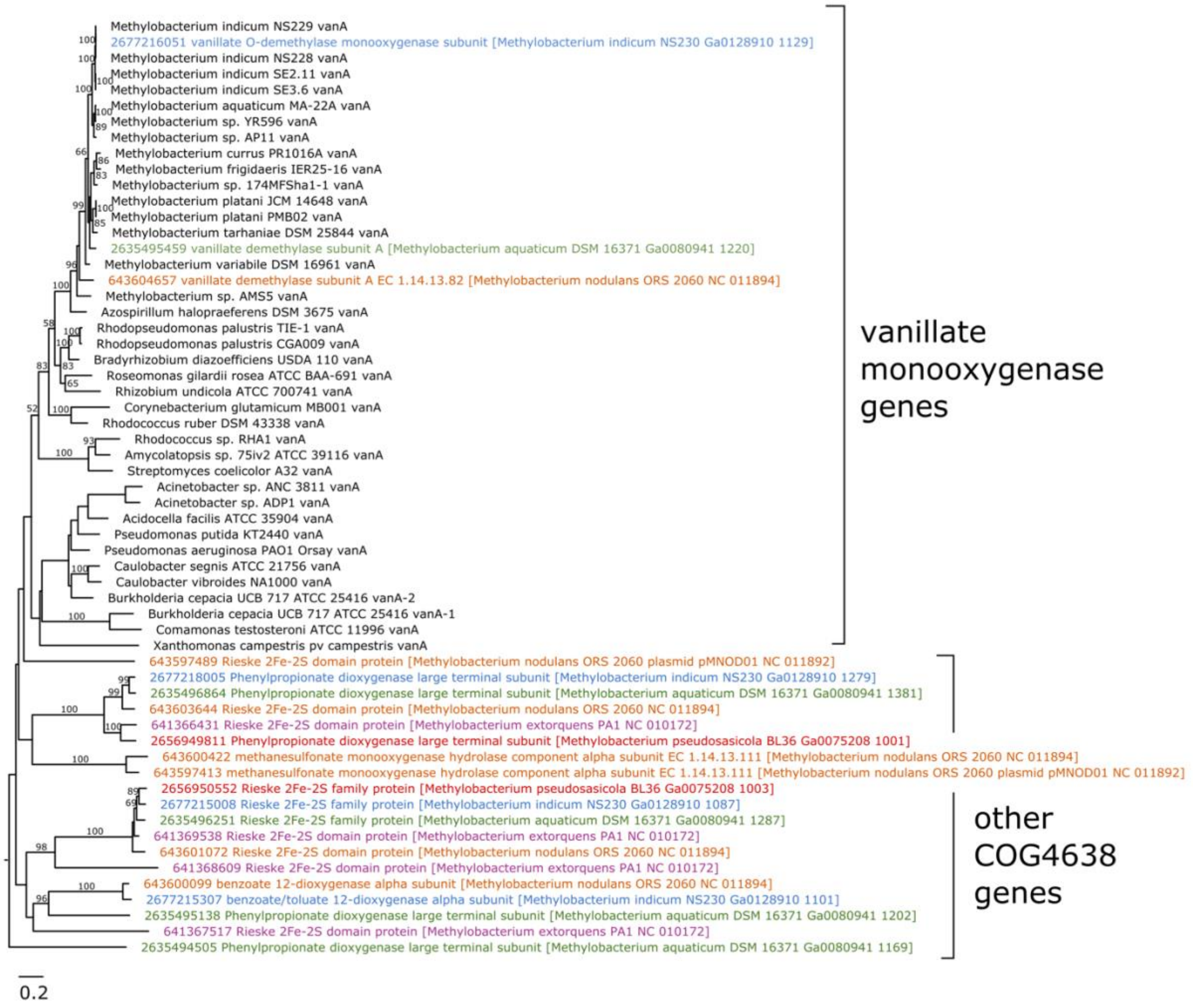


**Supplementary Figure 3.** Phylogeny of vanillate monooxygenase (*vanA*) genes with other COG 4638 (phenylpropionate dioxygenase or related ring-hydroxylating dioxygenase, large terminal subunit) genes from *M. nodulans*, *M. indicum* NS230, *M. aquaticum* DSM 16371, *M. pseudosasiacola*, and *M. extorquens* PA1. While several of the strains have multiple COG 4638 orthologs, only one from each organism (and none from *M. extorquens* or *M. pseudosasiacola*) clusters with known *vanA* genes. The gene alignment is provided in Supplementary Data Sheet 10.

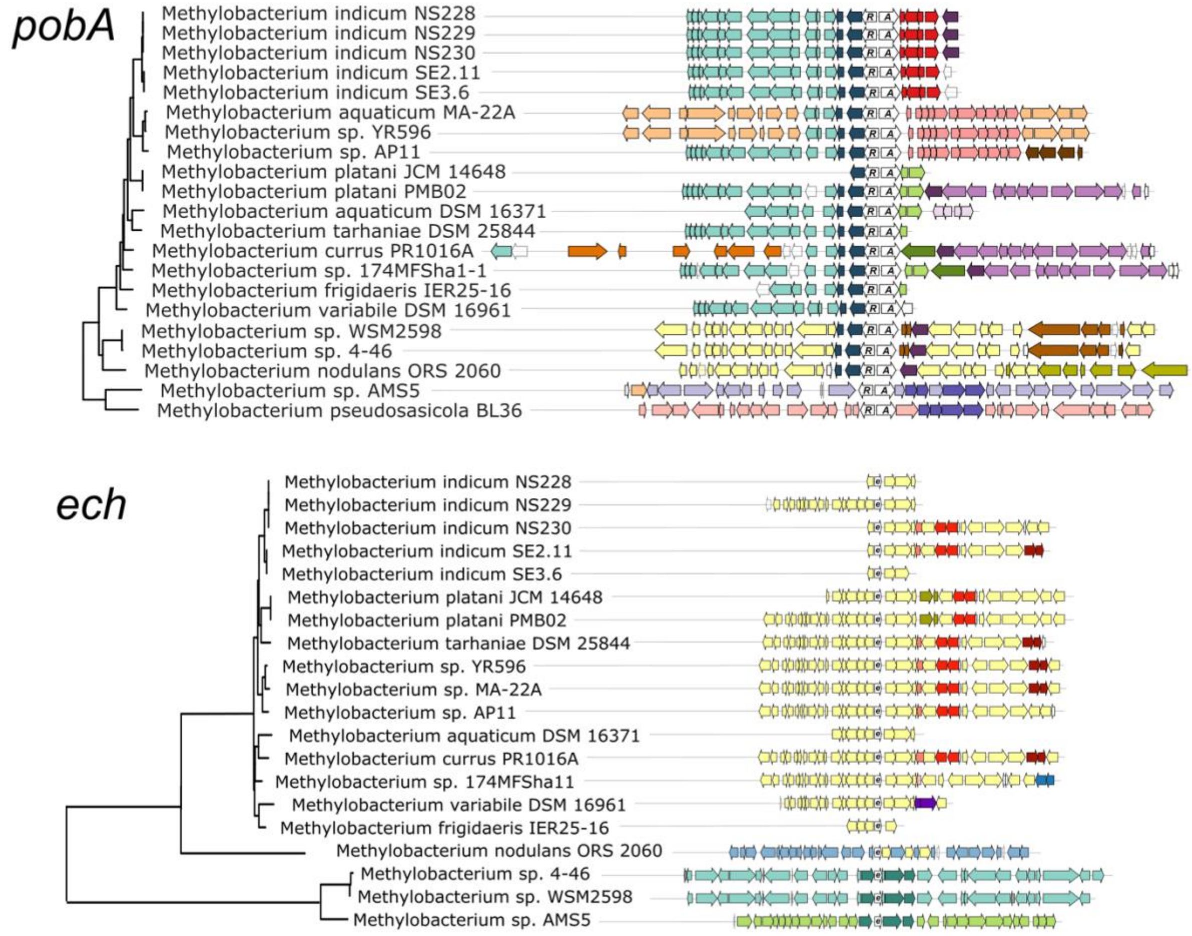


**Supplementary Figure 4.** Genomic context of aromatic catabolism genes *pobA* and *ech* in *Methylobacterium* species. As in Figure 3, genomic regions (from fully assembled genomes) or scaffolds (from partially assembled genomes) showing the neighborhoods of *pobA* and *ech* were aligned at the genes of interest. Gene phylogenies are excerpted from Fig. 2. Genes of interest and their operons are shown in white and labeled with initial letters. Within each of the two alignments, homologous regions shared across multiple genomes are shown in the same color to facilitate comparison among genomes.

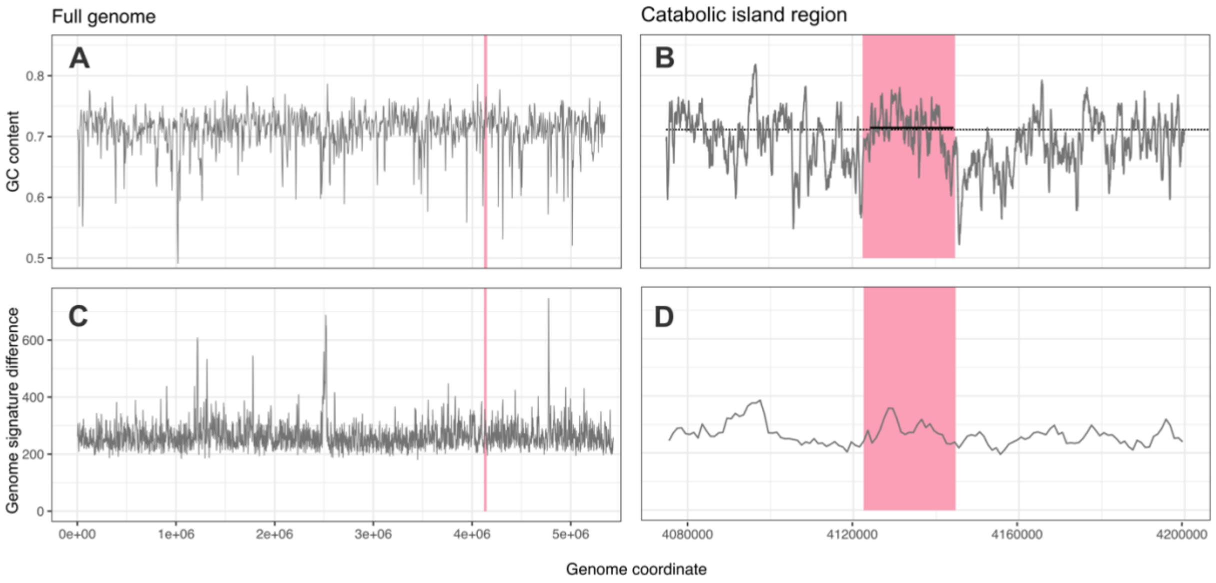


**Supplementary Figure 5.** *Methylobacterium sp*. AMS5 catabolic island has similar GC content and tetranucleotide composition to the rest of the genome. (A-B) GC content of the *Methylobacterium sp*. AMS5 genome. (A) The full genome, GC content calculated for a sliding 5 kb window. (B) The region surrounding the catabolic island, sliding 500 bp window; black dashed line shows average value for entire genome (0.711) and solid black line shows average value for catabolic island (0.714). (C-D) The genome signature difference index (Karlin, 2001) based on tetranucleotide abundances for a sliding 5 kb window, shown for (C) the full genome or (D) the region surrounding the catabolic island. High values indicate a large difference in tetranucleotide frequencies between that region and the genome average. In all plots, the region occupied by the catabolic island (20,028 bp) is highlighted in pink.

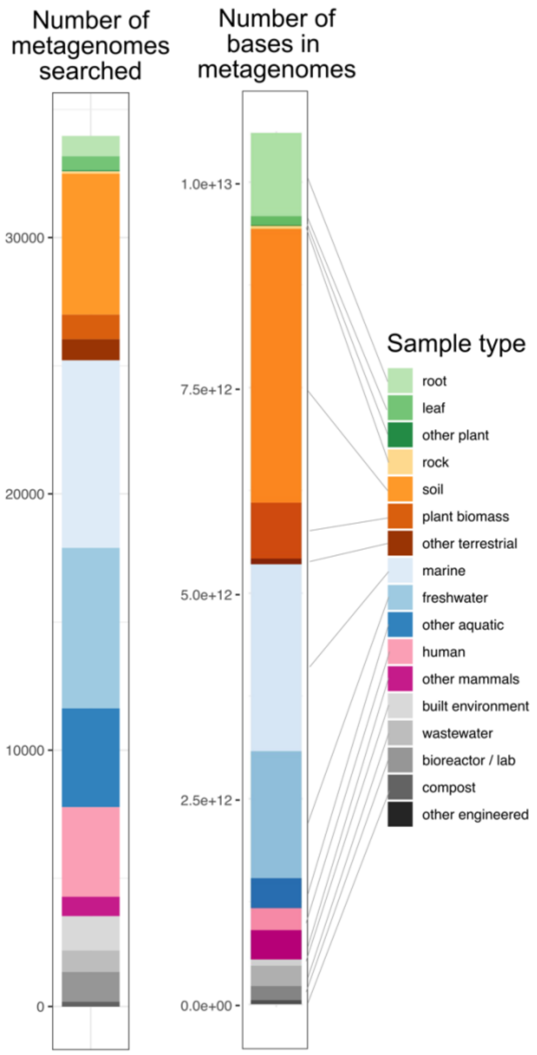


**Supplementary Figure 6**. Ecological distribution of metagenomes surveyed for *Methylobacterium* reads, for comparison with Fig. 8. "Number of bases in metagenomes" was calculated by summing the "Genome size, assembled" metric of the IMG metagenomes surveyed.

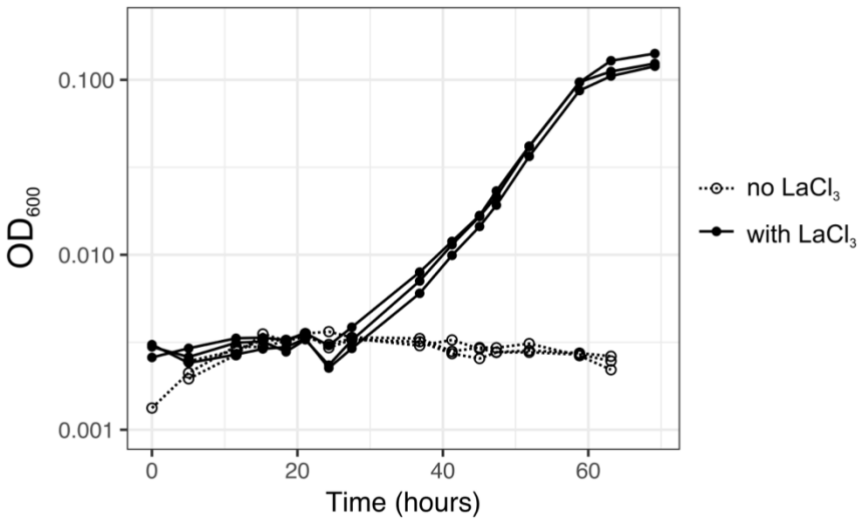


**Supplementary Figure 7**. *Methylobacterium* sp. 4-46 grows on methanol only when lanthanum is provided. Cultures were grown on MPIPES mineral medium with 15 mM methanol as the sole carbon source and either no $LaCl_3$ (open symbols and dotted lines) or 25 µM $LaCl_3$ (closed symbols and solid lines). Each line represents one biological replicate. Source data are given in Supplementary Table 12.
